# Astringent flavanol fires the locus-noradrenergic system, regulating neurobehavior and autonomic nerves

**DOI:** 10.1101/2025.05.06.652378

**Authors:** Yasuyuki Fujii, Shu Taira, Keisuke Shinoda, Yuki Yamato, Kazuki Sakata, Orie Muta, Yuta Osada, Ashiyu Ono, Toshiya Matsushita, Mizuki Azumi, Hitomi Shikano, Keiko Abe, Vittorio Calabrese, Naomi Osakabe

## Abstract

Astringent flavanols (FLs) have shown enhanced cognitive function in a large intervention trial, but their mechanism remains unclear due to poor bioavailability. An oral dose of FLs increased spontaneous locomotion and memory, with activation of the sympathetic-adrenergic-medullary (SAM) and hypothalamic-pituitary-adrenal (HPA) axis, as shown by increased urinary catecholamine concentrations and corticotropin-releasing hormone (CRH) mRNA expression in the paraventricular nucleus (PVN) of mice. MS imaging showed high intensity of noradrenaline (NA) in the locus coeruleus (LC), which plays a key role in learning and memory, the hypothalamus lateral preoptic area, which affects sleep and arousal, and the brainstem as the origin of the sympathetic nervous system, immediately after FLs administration. Subsequent in situ hybridization (ISH) analysis showed that these NAs originated from LC. Astringent compounds like FLs influence brain function and physiological changes. Furthermore, the present series of results presents novel mechanisms by which food sensory properties maintain homeostasis.

**Highlights:**

- A oral dose of astringent flavanols (FLs) immediately fires locus coeruleus-noradrenaline axis
- FLs enhance spontaneous locomotion and short-term memory in mice
- SAM and HPA axis are also activated by an oral dose of FLs
- FL’s astringency affects the brain and body, contributing to homeostasis.

## 1. INTRODUCTION

Astringency, a stimulant property unique to polyphenols, is present in only a few compounds. Flavanols (FLs), (-)-epicatechin and its oligomeric procyanidins (Fig. 1A) and anthocyanin are typical astringent compounds (Osakabe & Terao, 2018). These compounds are characterized by their high electrochemical activity and susceptibility to oxidative degradation (Cai et al., 2022). In particular, under neutral pH conditions, such as in the oral cavity or small intestine, they are known to immediately produce reactive oxygen species and decompose to produce decomposition products or oxides condensed with decomposition products(Friedman & Jürgens, 2000; Miller et al., 2008; Xue et al., 2024). FLs, a typical astringent substance, are abundant in cocoa, red wine, and berries. According to a systematic review of previous studies, FLs may play a role in maintaining or improving cognitive function (Andrews et al., 2023). Additionally, a recent large-scale intervention trial reported that one year of FLs intake restored hippocampal-dependent memory in older participants in the bottom third of habitual dietary quality with no habitual FLs intake (Brickman et al., 2023). In addition, several studies have reported that within approximately an hour of ingesting FLs, there were significant changes in brain function (Socci et al., 2017), e.g., increasing mental tracking capacity (Massee et al., 2015; Scholey et al., 2010), mood (Pandolfi et al., 2016), attention (Karabay et al., 2018), and restored working memory performance(Grassi et al., 2016). It has been suggested that these effects are accompanied by increased blood flow in the brain (Decroix et al., 2019; Francis et al., 2006) and peripheral circulation (Hooper et al., 2012; Sun et al., 2019).

**Fig. 1.**
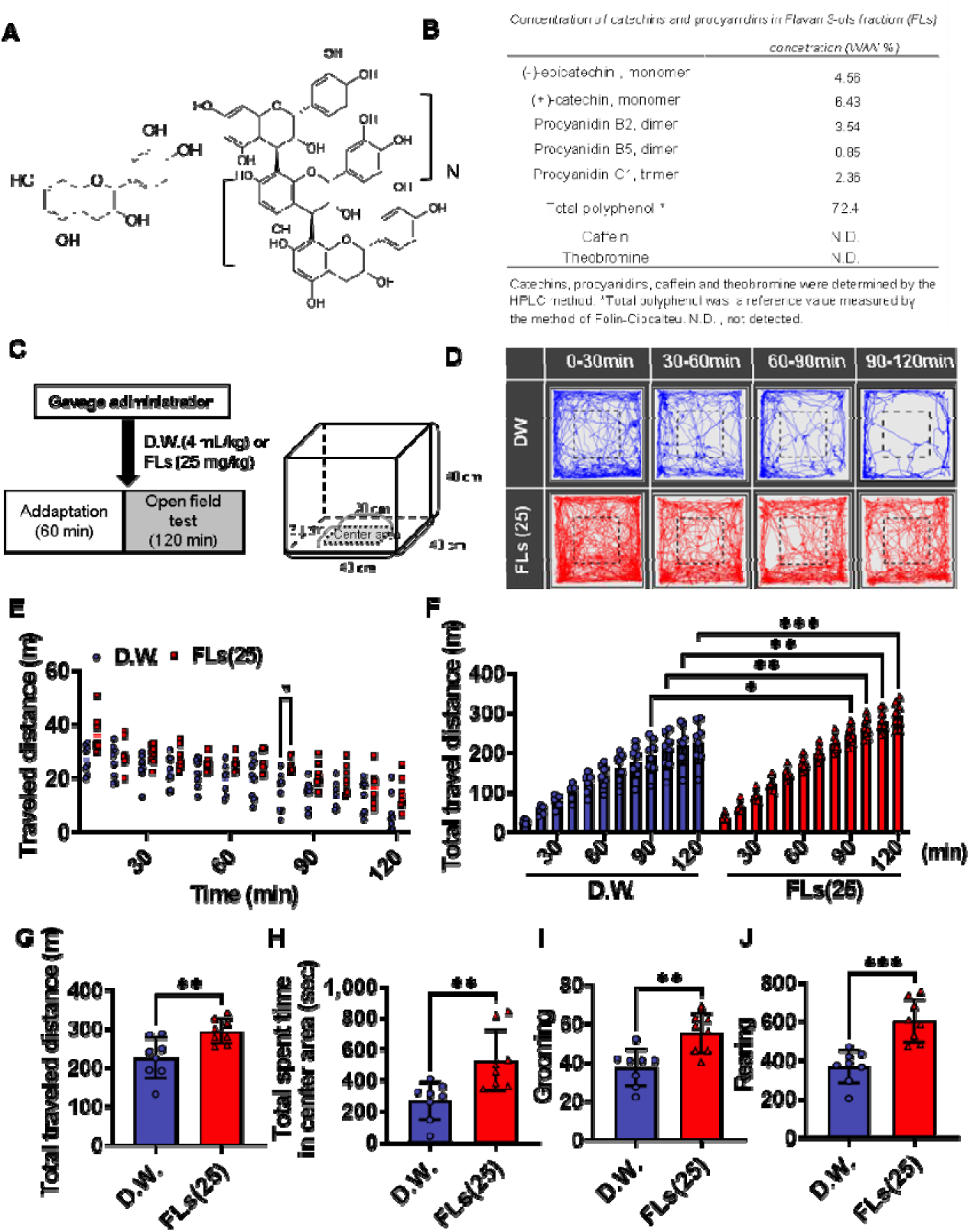
A single oral dose of flavanols (FLs) significantly increased mouse spontaneous activity. **A** chemical structure of flavanol;(-)epicatechin (left); procyanidins (right, n≥0). **B** The composition of FLs. Catechins, procyanidins, and xanthine derivatives were measured according to the method of Natsume et al.(Natsume et al., 2000). **C** Scheme of the open field test. The open-field test was conducted following the administration of D.W. (blue, n=8) or 25 mg/kg body weight flavanol (red, n=8). **D** The typical subsequent behavioral trajectories of mice in an arena over 120 min. **E** Distance traveled by each mouse every 10 mins. **F** Accumulated travel distance. **F** Total travel distance for 120 min. **G** The FLs-treated group significantly increased the total distance traveled compared with the D.W. group. **H** Duration spent in the center of the arena. **I** The number of grooming sessions. **J** The number of rearing sessions. The wakefulness indicators such as grooming and rearing were significantly increased in the FLs group compared to the DW group. To analyze multiple comparisons of travel distance or cumulative distance every 10 min, a two-way analysis of variance was performed, followed by Tukey’s multiple comparison test. A Student’s t-test was used to compare two groups of total travel distance, duration spent in the center of the arena, and the number of grooming and rearing sessions. The significance level is 0.05, with * p<0.05, ** p<0.01, and *** p<0.001.

Furthermore, meta-analyses have demonstrated an increase in peripheral flow-mediated dilatation (FMD) two hours after a dose of FLs (Sun et al., 2019). Our previous experimental animal studies found that this increase in blood flow after a single oral dose of FLs was due to hyperactivation of the sympathetic nervous systems (SNS) (Koizumi et al., 2021; Saito et al., 2016). However, FLs are known to have very poor bioavailability (Osakabe et al., 2022; Osakabe & Terao, 2018). It is unlikely that orally ingested FLs are distributed in blood and directly affect the central or autonomic nervous system, therefore, the mechanism by which this change occurs remains unknown.

This study aimed to investigate the mechanisms by which FLs promote brain function. To achieve this, we observed spontaneous locomotion and cognitive function in mice following a single oral administration of FLs. Furthermore, the impact of FLs on both stress response systems, on the sympathetic-adreno-medullar (SAM) and hypothalamic-pituitary-adrenal (HPA) axis was assessed by urinary catecholamine (CA) excretion and expression of corticotropin-releasing hormone (CRH) mRNA in the paraventricular nucleus (PVN) of the hypothalamus. In addition, we observed the changes in dynamics in the whole brain for L-dopa, dopamine (DA), noradrenaline (NA), and the metabolite normetanephrine (NMET) using the MS imaging technique and with the mRNA expression of the synthetic enzymes and transporters of these neurotransmitters were determined to be used in situ hybridization (ISH).

## 2. MATERIALS AND METHODS

### 2.1. Materials

The composition of FLs derived from cocoa used in the experiment as follows; (-)-epicatechin (monomer), 4.56%(W/W); (+)-catechin (monomer), 6.43%; procyanidin B2(dimer), 3.93%; procyanidin C1 (trimer), 2.36%; cinnamtannin A2 (tetramer), 1.45 % as shown in Fig.1B. Xanthine derivatives (caffeine, theobromine) were below the detection limit. All chemicals were measured according to the method of Natsume et al (Natsume et al., 2000).

### 2.2. Animals

Ten-weeks old male C57BL/6J mice were obtained from CLEA Japan, Inc. (Tokyo, Japan). During two weeks acclimation period, the animals were carefully handled to reduce anxiety behaviors. The study was conducted in accordance with the ARRIVE guidelines, and the protocol was approved by the Animal Experimentation Committee of Shibaura Institute of Technology (approval number: AEA 22007). As demonstrated in our previous research (Fujii et al., 2018), FLs have been shown to induce a stress response at doses of 10-50 mg/kg. Therefore, the dose used in the present experiment was set at 25 and/or 50 mg/kg.

### 2.3. Open filed test for observation of spontaneous behavior

We observed the effects of a single oral dose of FLs on the spontaneous behavior of mice in the open-field test. Mice were orally administrated with distilled water (D.W.), or FLs 25 mg/kg body weight. The open fields are composed of four transparent acrylic chambers aligned horizontally or vertically. Mice were immediately placed in the center of an arena (40 x 40 x 40 cm), and their behavior was recorded for 120 min (Fig.1C). The traveled distance was analyzed from the data, which were exported as a 30-min mp4 at 25 fps, corrected for tilt and brightness on filming using Premire Pro (Adobe Inc., California, USA), with a modification of the automatic tracking system developed by Zhang et al. (Zhang et al., 2020). All video files were processed by a MATLAB script running on a PC. The mouse activity code was designed to detect black mice such as C57BL/6J in a light arena.

### 2.4. Novel object recognition test

Novel object recognition test was performed according to the method of Ligar et al. with minor modifications as shown in Fig.2A (Leger et al., 2013). After two weeks of acclimation, mice were divided into the D.W.-treated group and the FLs 25mg/kg-treated group (n=8 each). A mouse given D.W. or FLs by gavage was placed in the center of an arena made from transparent acrylic (40 x 40 x 40 cm) and allowed to adapt for one hour. Two identical objects were placed in opposite quadrants of the arena (i.e., the northwest and southeast corners) and the mice were allowed 10 min to explore them freely. One hour later, one of the objects was replaced by a novel object of a different shape, and material, and the mice were allowed to explore the arena for 8 min to perform a novel object recognition test. The term “exploration” was defined as behavior exhibited by mice whereby the mouse’s noise is directed towards an object and within a distance of 2 cm from the object, with active vibrissae sweeping or sniffing. The tally did not include the time spent sitting on the object without indicating active exploration. T The decision exploring or not exploring was analyzed from the data, which were exported as a 30-minute mp4 at 25 fps, corrected for tilt and brightness on filming using Premire Pro (Adobe Inc., California, USA), with a modification of the automatic tracking system developed by Zhang et al. (Zhang et al., 2020). All video files are processed by a MATLAB script running on a PC. The mouse activity code was designed to detect black mice such as C57BL/6J in a light arena. To assess working memory, the discrimination index (DI) was calculated as the total exploration time to the novel object divided by the total exploration time, as previously described in a paper by Leger et al.(Leger et al., 2013) as follows. [(time spent exploring novel object) − (time exploring familiar object)]/[(time spent exploring novel object) + (time exploring familiar object)].

**Fig. 2.**
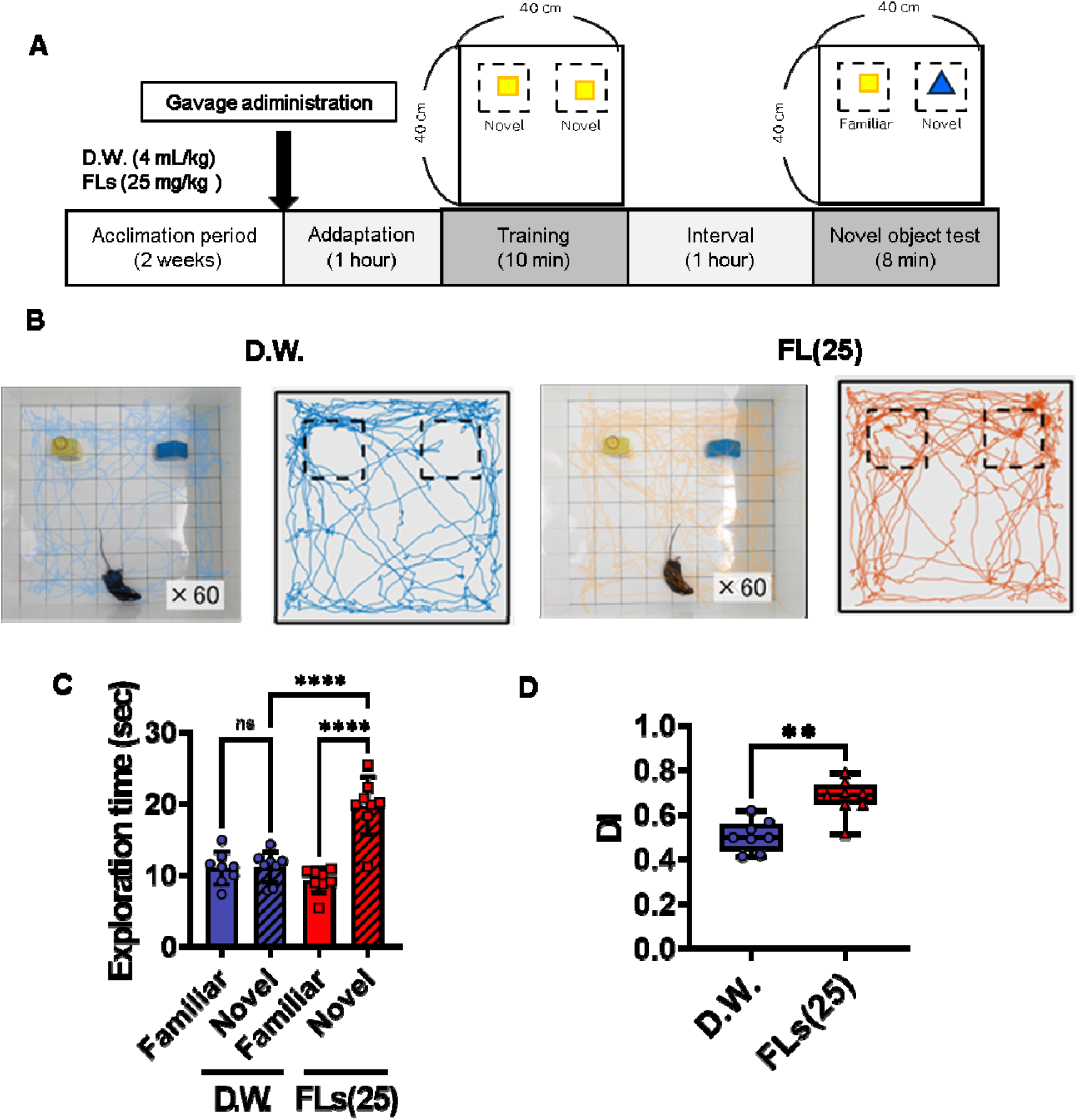
A single oral administration of flavanols (FLs) enhanced recognition memory in mice that was evaluated by a novel object test. **A** Scheme of the novel object test was conducted following the administration of D.W.(blue, n=8) or 25 mg/kg body weight flavanol (red, n=8) according to the method of Leger et al.(Leger et al., 2013). **B** The typical subsequent trajectories by mice during the 8 min following exposure to a novel object (right, D.W.; left, FLs 25 mg/kg). **C** Total exploration time for each group of objects. **D** The discrimination index (DI) results were expressed as a percentage of the total exploration time divided by the total exploration time to the novel object. To analyze multiple comparisons of exploration time, a two-way analysis of variance was performed, followed by Tukey’s multiple comparison test. For comparisons between two groups of DI, the Mann-Whitney test was used. The significance level was 0.05, with ** p<0.01 and **** p<0.0001.

### 2.5. Quantification of urinary CA excretion using HPLC

It has been suggested that brief periods of social isolation in metabolic cages can markedly alter SNS activity in mice (Takahashi, 2022). In our preceding research, we demonstrated that co-housing two mice in metabolic cages markedly reduced the stress response associated with single housing (Muta et al., 2023). Consequently, in the present study, we employed this approach to examine the SNS activity of FLs. Following a 48-hour acclimation period, urine was collected for 24 hours using a tube containing 20 µl of 2.5 mol/L HCl following oral administration of test chemicals (Fig.3A). Oral administration of DW or FLs was performed between 10:00 and 11:00.

**Fig. 3.**
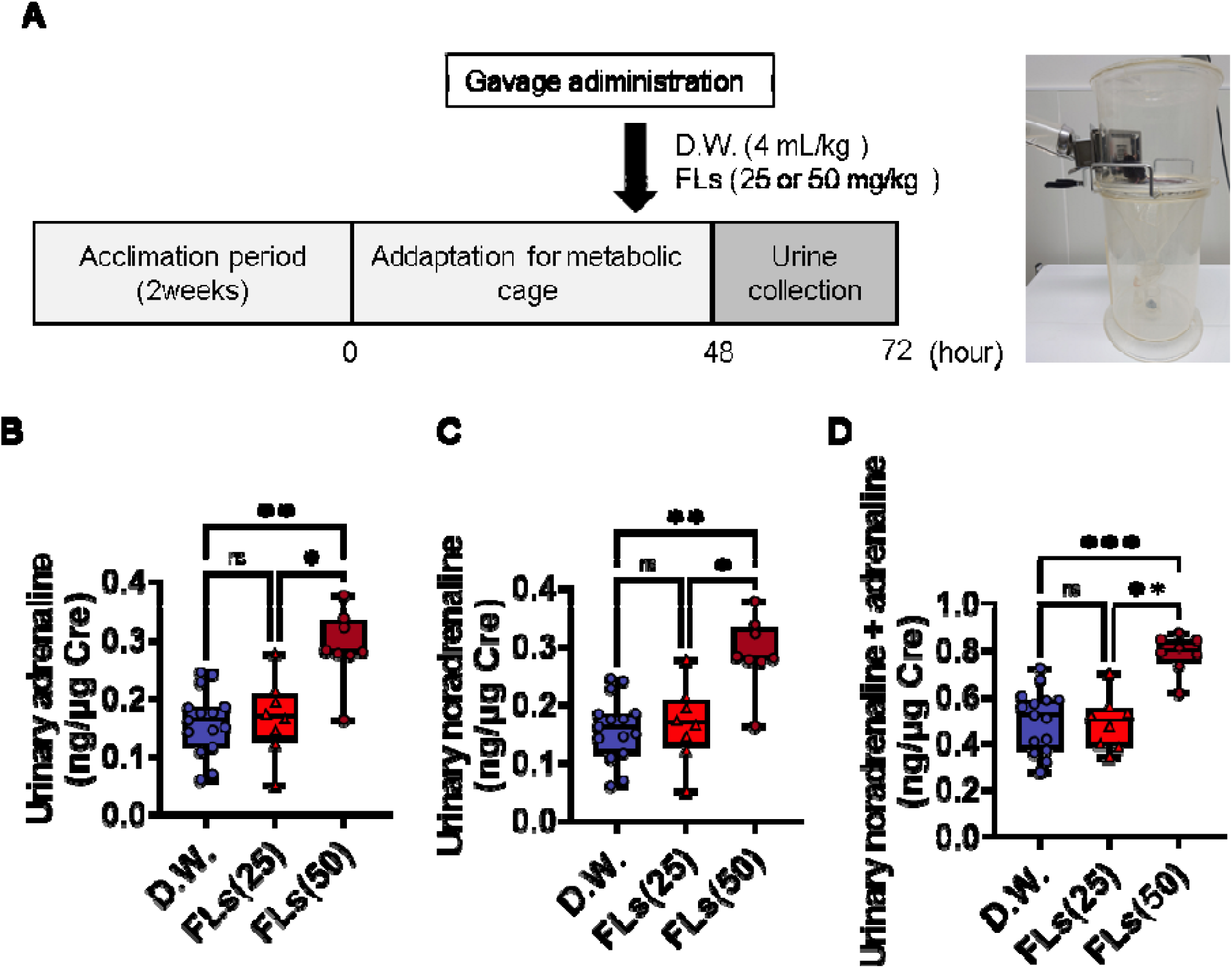
A single oral dose of flavanols (FLs) increased urinary excretion of CA. **A** Scheme for administration and urine collection protocol. After a 48-hrs acclimation period, 24-h urine samples were collected according to the method of Muta et al. (Muta et al., 2023). Urinary excretion of **B** noradrenaline (NA), **C** adrenaline (AD), and **D** total CA analysis in 24 hrs-urine (D.W. in blue, n=15), 25 mg/kg (in light red) or 50 mg/kg (in dark red) flavanol (FLs, n=8 pair each). CA excretion was expressed as a ratio to the urinary creatinine concentration. Statistical analysis for CA concentration was performed using the Kruskal-Wallis test followed by Dunn’s multiple comparisons test. The significance level was 0.05, * p<0.05, ** p<0.01 and *** p<0.001.

NA and adrenaline (AD) were purchased from Sigma–Aldrich. The urine was neutralized and heated for 10 min following incubation with 500 U/ml of the enzyme sulfatase from Helix pomatia Type H-2 (Sigma–Aldrich) for one hour at 37°C. Following this, isoprenaline (Sigma–Aldrich) was added as the internal standard, after which CA was purified using Monospin PBA® (GL Sciences, Tokyo, Japan). The HPLC system (Prominance HPLC System Shimazu Corporation, Kyoto, Japan) comprised a quaternary pump with a vacuum degasser, a thermostatted column compartment, and an autosampler equipped with an electrochemical detector (ECD 700 S; Eicom Corporation, Kyoto, Japan). A reverse-phase column (Inertsil ODS-4, 250 × 3.0 mm ID, 5 μm; GL Sciences) was utilized, with the column temperature maintained at 35°C. The HPLC mobile phase comprised 24 mM acetate-citrate buffer (pH 3.5, -CH3CN, 100/14.1 v/v). The mobile phase flow rate was 0.3 ml/min, with an injection volume of 20 μl. Eluents were detected and analyzed at 500 mV. The excretion of CA was expressed as a ratio with the urinary creatinine concentration, which was measured using Laboassay creatinine (FUJIFILM Wako Pure Chemical Corporation).

### 2.6. Observation of c-fos and CRH dynamics using ISH

Mice were gavaged with either D.W. or 25 mg/kg FLs and then decapitated 15, 30 or 60 min later (Fig.4A). The excised brains were immediately frozen and coronally sectioned (8 μm thick) using a cryostat (Leica, Wetzlar, Germany). The sections of fresh-frozen brains were prepared with a cryostat and thaw-mounted on glass slides (Matsunami Glass Ind. Ltd., Osaka, Japan). ISH was carried out to evaluate c-fos and CRH mRNA expression with the RNAscope® Multiprex Assay (Advanced Cell Diagnostics, CA, USA). An ImmEdge™ pen (H-4000, Vector Laboratories Inc., California, USA) was used to create a barrier around the sections. The barriers were dried completely at room temperature. The samples were treated with hydrogen peroxide (RNAscope® H_2_O_2_ and Protease Plus Reagents, Advanced Cell Diagnostics, CA, USA) for 10□min at room temperature, and washed twice in D.W. The antigens were activated with RNAscope® Target Retrieval Reagents (Advanced Cell Diagnostics) for 15□min, washed immediately in D.W., and dehydrated in ethanol for 1□min. The sections were treated with Protease Plus Reagents for 30□min at 40 °C in the HybEZ™ OVEN (Advanced Cell Diagnostics); then the sections were washed twice in D.W. Next, the RNA probe solution was mixed (1:50 dilution) with the target RNA probes: Mm-Fos (316921, Advanced Cell Diagnostics) and Mm-Crh-C2 (316091-C2, Advanced Cell Diagnostics). The sections were incubated with the probes for 2□h at 40□°C, then washed twice in wash buffer (Advanced Cell Diagnostics, CA, USA). Next, the sections were treated serially with AMP1 for 30□min (Advanced Cell Diagnostics), AMP2 for 15□min (Advanced Cell Diagnostics), AMP3 for 30□min (Advanced Cell Diagnostics), AMP4 for 15□min (Advanced Cell Diagnostics) at 40□°C, and AMP5 for 30□min (Advanced Cell Diagnostics), and AMP6 for 15□min (Advanced Cell Diagnostics) at room temperature. The sections were washed twice in the wash buffer for 2□min and treated with red conjugates (60:1), Fast red A and Fast red B, for 10□min at room temperature. After washing in wash buffer, the sections were treated successively with AMP7 for 15□min (Advanced Cell Diagnostics), AMP8 for 30□min (Advanced Cell Diagnostics) at 40□°C, and AMP9 for 30□min (Advanced Cell Diagnostics), and AMP10 for 15□min (Advanced Cell Diagnostics) at room temperature. Next, the sections were treated with green conjugates (50:1) as Fast green A and Fast green B (Advanced Cell Diagnostics), for 10□min at room temperature. The sections were rinsed twice in D.W. Finally, the sections were immersed in 50% Gill hematoxylin (GHS116-500ML, Sigma Aldrich, St-Louis, USA) for 30□s and washed twice immediately in water. After drying completely in a dry oven (MIR-162-PJ, Panasonic, Osaka, Japan) at 60□°C, the sections were covered with Vector mount™ permanent mounting medium (H-5000, Vector Laboratories Inc., California, USA).

**Fig. 4.**
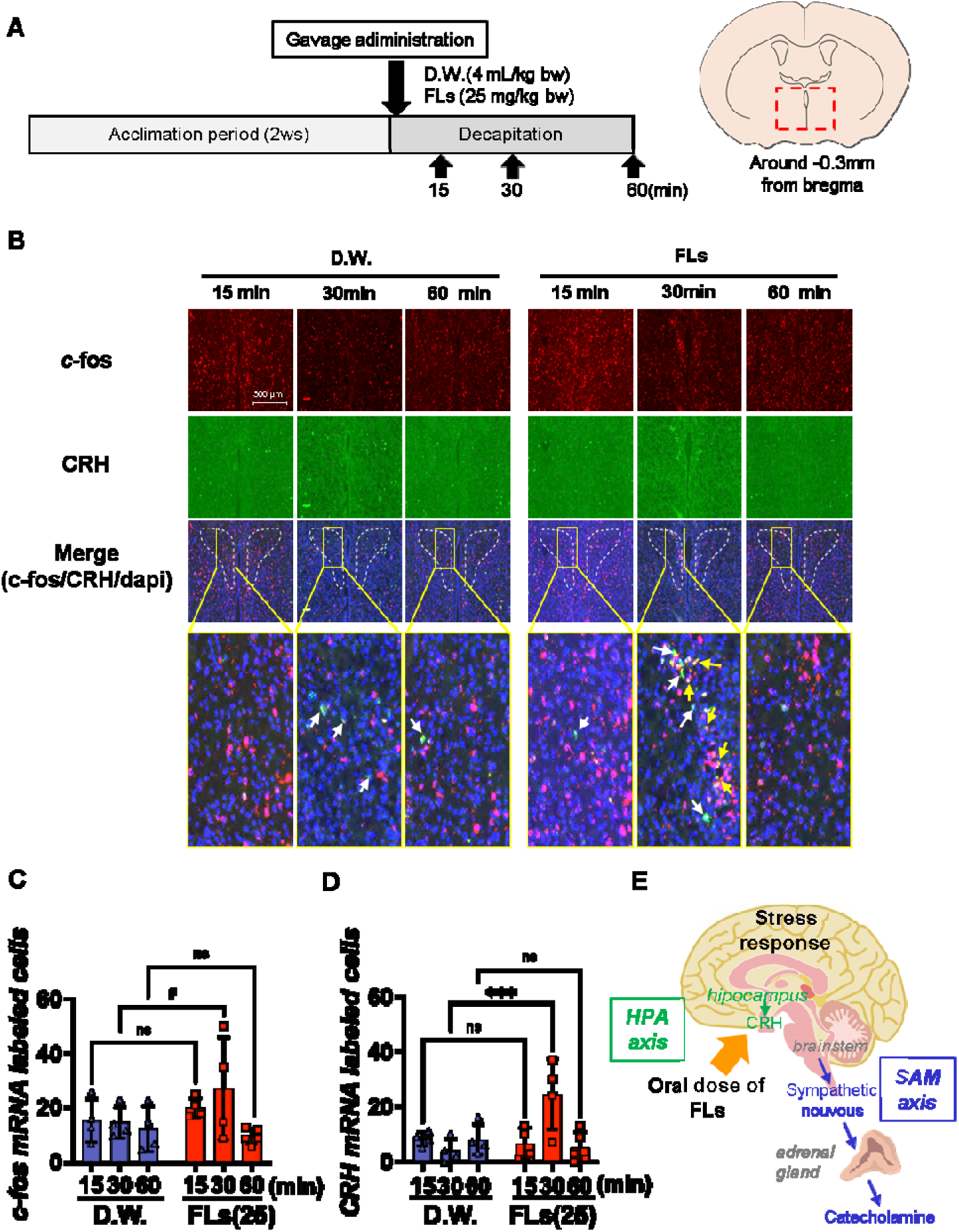
A single oral administration of flavanols (FLs) induced corticotropin-releasing hormone (CRH) mRNA in the paraventricular nucleus (PVN) of the hypothalamus. **A** Scheme of administration and brain collection protocol according to the method of Fujii et al. (Fujii et al., 2018). **B** A typical c-fos (top) and CRH mRNA (center) detected by ISH, as well as their merged and enlarged images (bottom, green as cfos; pink, CRH; blue, dapi). mRNA expression of cfos was demonstrated by the white arrow, and its CRH was shown by the yellow arrow in PVN. **C** The number of c-fos-positive cells, and **D** the number of CRH-positive cells in PVN enclosed by dotted line (n = 4 to 5). Compared to the D.W.-treated group, the number of CRH mRNA-expressing cells significantly increased in the FLs-treated group 30 min after administration.. To analyze multiple comparisons of the number of positive cells, a two-way analysis of variance was performed, followed by Tukey’s multiple comparison test. The significance level was 0.05, with *** p<0.001. **E** A diagram of a single oral dose of FLs enhanced the stress response system. SAM, sympathetic-adrenergic-medullary axis; HPA, hypothalamic-pituitary-adrenal axis.

### 2.7. Observation of noradrenaline dynamics using MS imaging and ISH

Mice were gavaged with either D.W. or 25 mg/kg FLs and then decapitated immediately after, 15 or 60 min later(Fig.5A). The excised brains were immediately frozen and sectioned (8 μm thick) using a cryostat (Leica, Wetzlar, Germany). The sections of fresh-frozen brains were prepared with a cryostat and thaw-mounted on conductive indium-tin-oxide-coated glass slides (Matsunami Glass Ind. Ltd., Osaka, Japan). A pyrylium-based derivatization method was applied for the tissue localization imaging of NA, its precursors L-dopa and DA, and its metabolite NMET according to the method by previous publication (Shikano et al., 2022). In brief, a solution of TMPy (4.8 mg/200 µL) (Taiyo Nippon Sanso Co., Tokyo, Japan) was applied to brain sections using an airbrush (Procon Boy FWA Platinum 0.2-mm caliber airbrush, Mr. Hobby, Tokyo, Japan). To enhance the reaction efficiency of TMPy on sections, the TMPy-sprayed sections were placed into a dedicated container and allowed to react at 60 °C for 10 min. The container contained two channels in the central partition, to wick moisture from the wet filter paper region to the sample section region. The filter paper was soaked with 1 mL methanol/water (70/30 volume/volume) and placed next to the section inside the container, which was then completely sealed to maintain humidity levels. The TMPy-labeled brain sections were sprayed with matrix (α-cyano-4-hydroxycinnamic acid-methanol/water/TFA = 70/29.9/0.1 volume/volume) using an automated pneumatic sprayer (TM-Sprayer, HTX Tech., Chapel Hill, NC, USA). Ten passes were sprayed according to the following conditions: flow rate, 120 μL/min; airflow, 10 psi; nozzle speed, 1100 mm/min. Each section was scanned to detect the laser spot area, and laser spot areas (200 shots) were detected with a spot-to-spot center distance of 120 µm using MALDI-Q-TOF MSI (rapifleX®, Bruker Daltonics Bremen, GmbH). The section surface was irradiated with yttrium aluminum garnet laser shots in the positive ion detection mode. The laser power was optimized to minimize the in-source decay of targets. Signals between m/z 200-800 were collected. Obtained mass spectrometry spectra were reconstructed to produce mass spectrometry images using Scils Lab software (Bruker Daltonics). Optical images of brain sections were obtained using a scanner (GT-X830, Epson, Tokyo, Japan) followed by MALDI-TOF MS of the sections. The detected masses of TMPy-labeled standard chemicals increased by 105.0 Da compared with the original mass, TMPy-l L-dopa, m/z 302.1; TMPy-l DA, m/z 258.1; TMPy-l NA, m/z 274.1 ; TMPy-l NMET, m/z 288.2. Tandem mass spectrometry confirmed the fragmentation ions of TMPy from the standard sample. A fragmented ion of the pyridine ring moiety was regularly cleaved and observed for all TMPy-modified target molecules.Tandem mass spectrometry confirmed the fragmentation ions of TMPy from the standard sample.

**Fig. 5.**
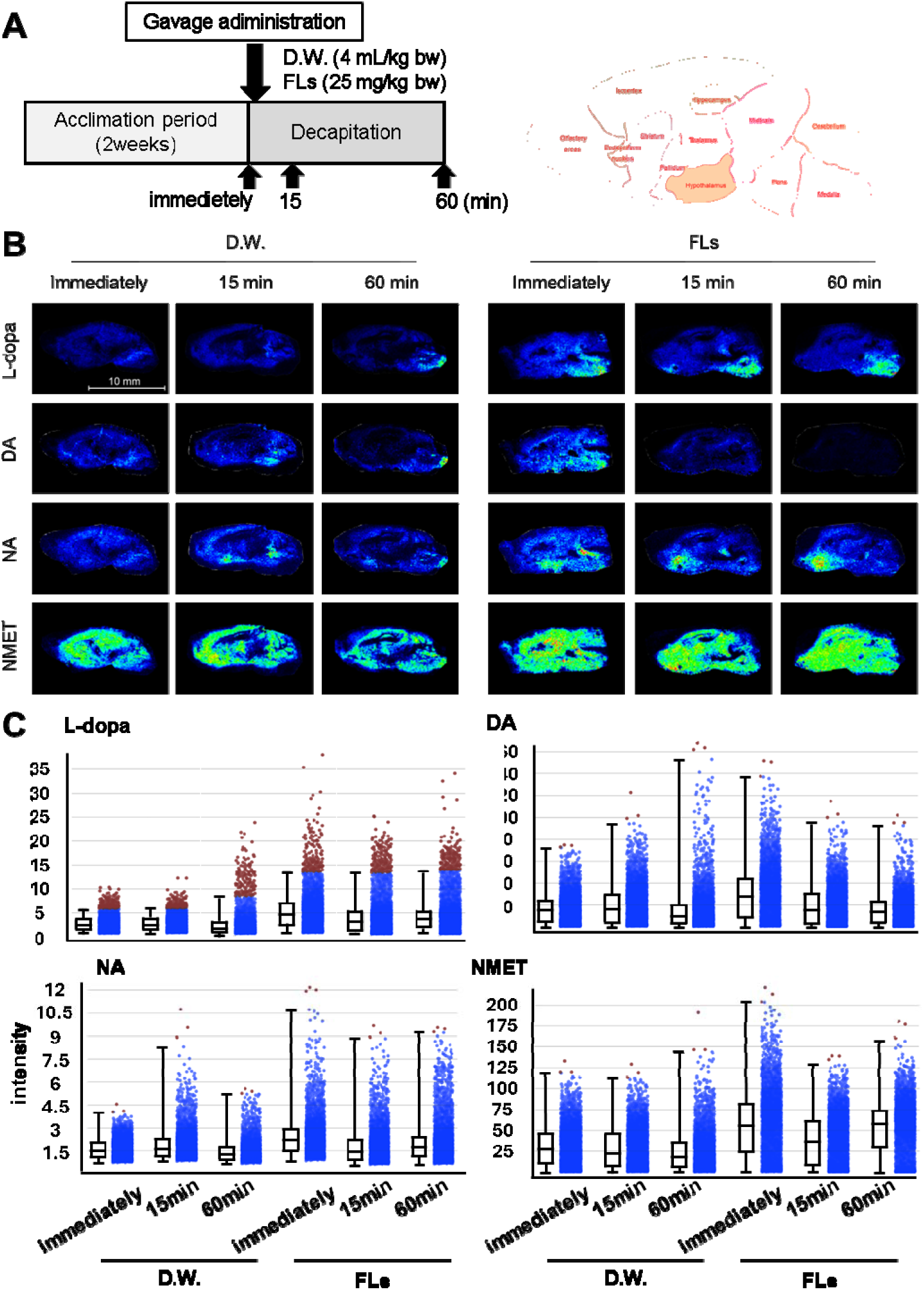
FLs caused substantial perturbations in L-dopa, dopamine, and noradrenaline FL and the metabolite noretinephrine (NMET) in the whole mouse brain. A Diagram of the dosing and brain harvesting protocol and a schematic of the mouse brain. Mice were given a single oral dose of D.W. or 25 mg/kg flavanols, brains harvested by decapitating immediately, and 15 or 60 min after administration. Excised brains were immediately frozen and sectioned (8 μm thick) using a cryostat. B Representative MS images of L-dopa (top), dopamine (DA, second from top), noradrenaline (NA, third from top), and normetanephrine (NMET, bottom). C Relative intensity of L-dopa, DA, NA, and NMET of the image B.

### 2.8. Fluorescence microscopy and quantification

All image analyses were performed by experimenters blinded to the experimental conditions. ISH image was taken using a fluorescence digital microscope (BZ-X800, KEYENCE Corp., Osaka, Japan). Cell counting of c-fos and CRH were performed throughout the extend of PVN. Each expression was quantified using NIH Image J software ver. 1.52 (http://rsb.info.nih.gov/ij/index.html).

### 2.9. Statistical Analysis

The sample size was determined from the results of a preliminary experiment using a power test with a significance level of 0.05 and power 0.9. All quantitative assessments were carried out under blinded manner. No data was excluded from the analysis. All data were expressed as mean ± standard deviation. All data were analyzed using GraphPad Prism 10.2.3 (San Diego, CA, USA). The normal distribution of the sample was tested using the Shapiro-Wilk test. Comparisons between the two groups were made using the Student’s t test or the Mann-Whitney test. For statistics on urinary CAs, the Kruskal-Wallis test was performed followed by Dunn’s multiple comparisons test. Otherwise, two-way ANOVA followed by the Tukey’s multiple comparisons test as post hoc test was used for multiple comparisons. The significance levels were defined at # p<0.1, * p<0.05, ** p<0.01, *** p <0.001.

## 3. RESULTS

### 3.1. Astringent FLs increased spontaneous activity

The 120 min open field results after intragastric administration of D.W. or 25 mg/kg FLs were shown in the line plot in Fig. 1D, and the distance traveled every 10 min in Fig.1E. The cumulative distance traveled by the FLs-treated group was significantly longer than that of the DW-treated group, from 90 to 120 min after administration (Fig. 1F). The total distance traveled (Fig.1G), time spent in the central area (Fig.1H), number of grooming (Fig. 1I), and rearing sessions (Fig. 1J) were all significantly higher in the FLs group than in the DW group.

### 3.2. Astringent FLs enhanced cognitive function in novel object recognition test

Fig. 2B showed the line plot results of mice in the novel object test after a single dose of D.W. (left) or FLs (right). Typical mouse behavior was also shown in the supplemental video(SVideo1 and 2). Fig. 2C displayed the total search time for the two objects, while Fig. 2D showed the Discrimination Index (DI). The FLs treatment group showed a significant increase in search time for the novel object (Fig. 2C), which led to a considerable rise in DI (Fig.2D).

### 3.3. FLs increased urinary CA excretion by activating SAM axis

Fig. 3 illustrates the impact of a single dose of FLs on SNS activity by quantified urinary CA excretion over 24 hr. The total urinary noradrenaline (Fig. 3B), adrenaline (Fig. 3C), and total CA (Fig. 3D) levels demonstrated a notable elevation in the 50 mg/kg FLs group, in comparison to the D.W. group, although this was not observed in the 25 mg/kg FLs group.

### 3.4. FLs increased mRNA expression of c-fos and CRH in PVN by activating HPA axis

Fig. 4 shows the results of observing the mRNA expression of cfos and the stress hormone CRH in the hypothalamic PVN using the ISH method. The upper part of Fig. 4B shows c-fos, the middle part shows CRH, and the lower part shows the merged images of c-fos, CRH, and dapi. he white or yellow arrow indicated cfos or CRH mRNA signal. The number of cells expressing c-fos (Fig. 4C) and CRH (Fig. 4D) in the PVN enclosed by the dotted line was counted using these images. There was a slight increase in the number of cfos-expressing cells between the D.W. group and the FLs group 30 min after FLs administration (Fig. 4C). CRH-expressing cell number showed a significant increase in expression 30 min after FLs treatment (Fig. 4D). The findings indicate that a single oral dose of FLs elicits a stress response and stimulates the SAM and HPA axes (Fig. 4E).

### 3.5. FLs exerted a substantial impact on the dynamics of noradrenaline in the whole mouse brain

Fig. 5B showed the distribution of NA (third from the top), its precursor 3-hydroxy-L-tyrosine (L-dopa, top) and DA (second from top), and its metabolite NMET (bottom), detected in sagittal sections of mouse brain immediately after, 15 or 60 min after a single dose of D.W. (left) or 25 mg/kg FLs (right). L-dopa and NMET were detected at higher levels in the FLs group than D.W. group throughout the observation period. A higher intensity of DA was observed in the hypothalamus, midbrain, pons, and medulla immediately after administration of FLs, and was rarely detectable 15 or 60 min after administration. It was also observed a higher level of NA in parts of the hypothalamus, pons, and medulla immediately after administration of FLs. In addition, a greater NA was observed in the ventral striatum 15 or 60 min after administration of FLs. The respective intensities of L-dopa, DA, NA, and NMET were shown in Fig. 5C after administration of D.W. or FLs. The intensity of L-dopa and NMET was higher in the whole brain throughout the observation period by a single oral dose of FLs. DA intensity was greater immediately after the administration of FLs. Immediately or 60 min after a dose of FLs, the intensity of NA was elevated.

### 3.6. FLs substantially increased noradrenaline intensity within LC, lateral preoptic area (LPO), and nucleus accumbens (NAc)

In Fig.6, the distribution and relative intensity of NA in LC, LPO and NAc. Fig.6A shows the change in NA in LC after administration of FLs. Immediately after the administration of FLs, the intensity of NA in the LC markedly increased. Fig. 6B showed the change in NA in LPO after administration of FLs. The NA in LPO observed higher intensity at all durations in the FLs-administered group compared to the D.W.-treatment group. In Fig.6C, NA signals in NAc following treatment of D.W. or FLs. The intensity of NA increased over time with FL administration.

**Fig. 6.**
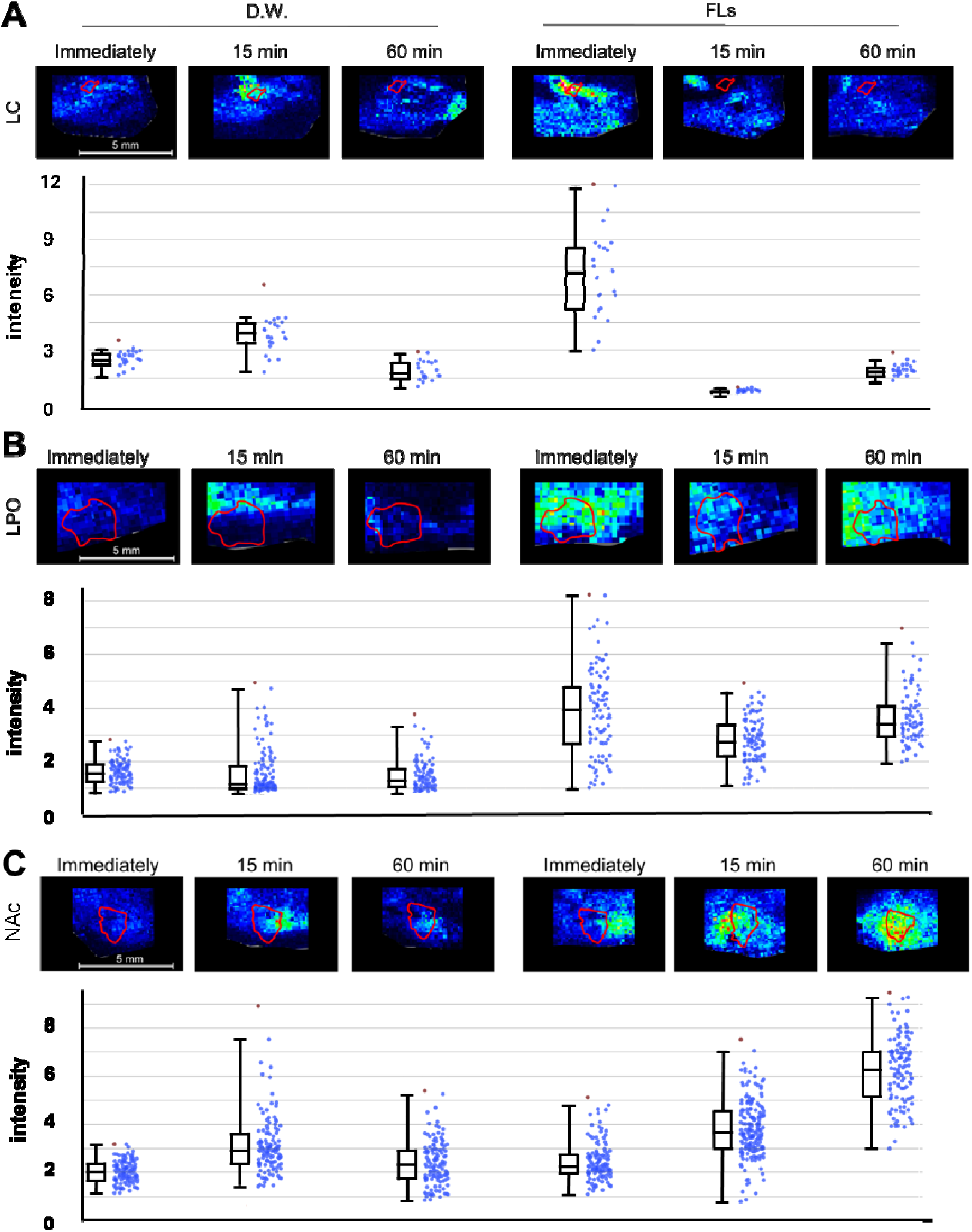
A single oral administration of flavanols (FLs) immediately enhanced the intensity of noradrenaline (NA) in locus coeruleus (LC), lateral preoptic area (LPO) and nucleus accumbens (NAc). **A** Representative MS images and relative intensity of NA in LC. **B** Representative MS images and relative intensity of NA in LPO. **C** A Representative MS images and relative intensity of NA in NAc.

### 3.7. FLs resulted in an augmentation of noradrenaline biosynthesis or transporter production enzymes within the midbrain

Given that a single oral administration of FLs resulted in high localized NA intensity, we analyzed the expression of enzymes involved in noradrenaline biosynthesis (Fig.7A) or transporter production. Fig. 7B shows the mRNA expression of each enzyme and its enlarged image in frozen brain serial sections obtained immediately after FLs administration, and 15 and 60 min later using the same method as in Fig. 6. From the top to the bottom, the biosynthetic pathways (Fig. 7A), tyrosine hydroxylase (TH), dopamine β-hydroxylase (DBH), as well as a transporter, vesicular monoamine transporter (VMAT)2 were shown marged with dapi. In the FL group, as shown by the yellow arrows, a greater increase in TH mRNA expression was observed in the LC (yellow arrow) and ventral tegmental area (VTA, white arrow) immediately after administration, but it was attenuated after 15 or 60 min. On the other hand, in the D.W. group, it was only slightly detected 15 min after administration. DBH mRNA expression was observed as a strong signal in the LC (yellow) immediately after FLs administration. On the other hand, in the D.W. group, it was only slightly detected 15 min after administration. In addition, VMAT2 was marked increased in the LC (yellow) and VTA (white) immediately after administration in the FL group, and only slightly observed 15 min after administration in the D.W. group.

**Fig. 7.**
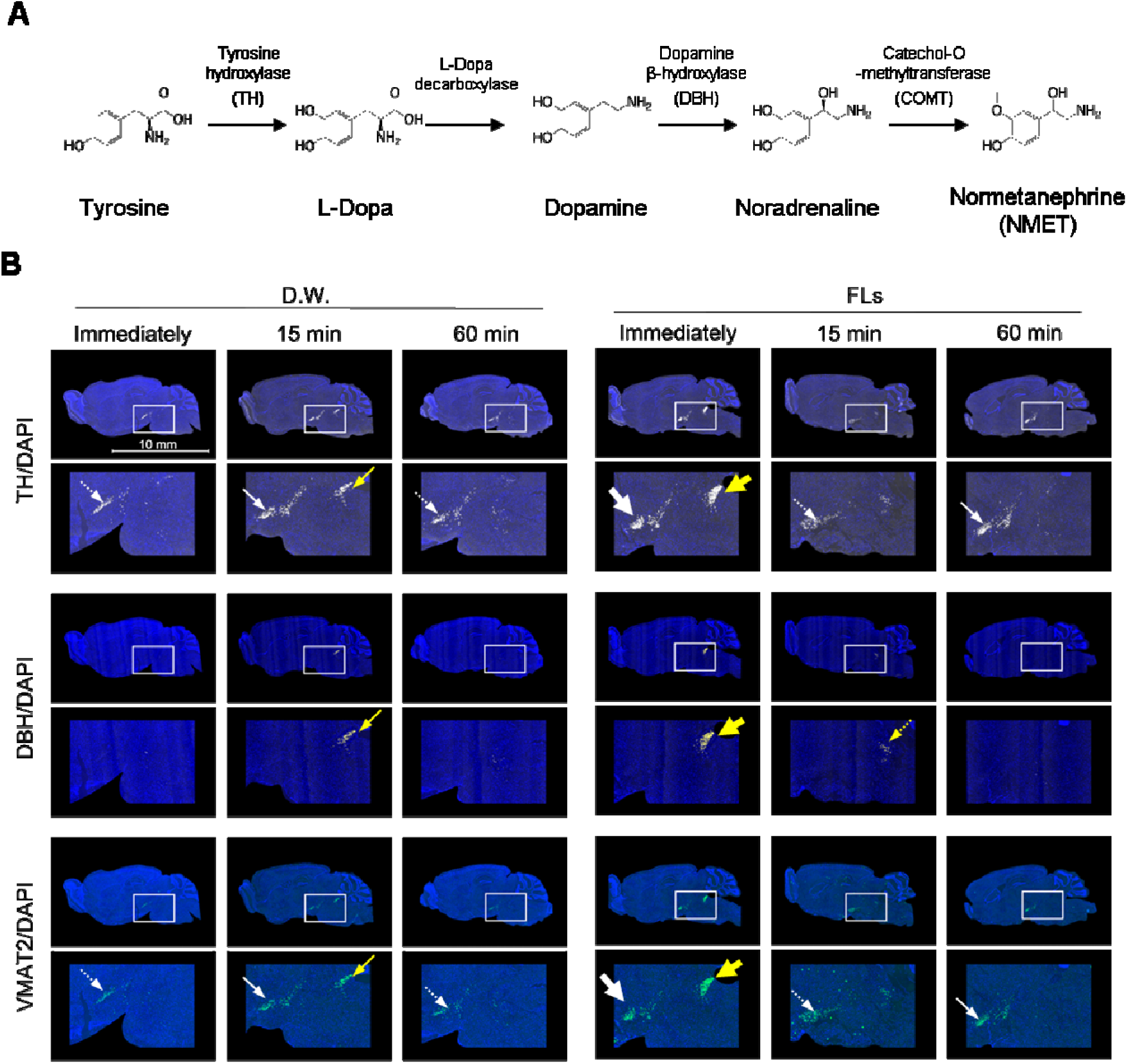
A single oral administration of flavanols (FLs) induced mRNA of synthesis enzyme and transporter for noradrenaline (NA) or dopamine (DA) in locus coeruleus (LC) or ventral tegmental area (VTA). **A** Pathways for the metabolism of DA and NA. **B** The mRNA expression of tyrsine hydroxylase (TH, top), dopamine β-hydroxylase (DBH, second from top), and vesicular monoamine transporter 2 (VMAT2, bottom) was assessed by using the ISH method. The expression in the LC was demonstrated by yellow arrows, and the expression in the VTA was shown by white arrows.

## 4. DISCUSSION

The series of experiments revealed that a single oral administration of FLs had a marked impact on neurotransmitter dynamics throughout the brain. In particular, the substantial increase in NA in LC and the induction of mRNA expression of the synthesizing enzyme and transporter in LC indicate the activation of the noradrenergic neuronal network (Fig. 5-7).

LC is the source of NA in the forebrain and is projected throughout the forebrain, brainstem, cerebellum, and spinal cord (Breton-Provencher et al., 2021). Furthermore, most cortical and subcortical regions are known to be under the control of LC-NA axonal nerves (Nomura et al., 2014; Zerbi et al., 2019). It is established that the release of NA from LC neurons plays a role in regulating wakefulness. Optogenetic techniques have revealed frequency-dependent correlations between local neuronal firing, cortical activity, sleep-to-wake transition, and general motor arousal in the LC of the mouse (Carter et al., 2010). Conversely, during the sleep cycle, there is a reduction in systemic LC-NA activity. However, LC spikes generated by sensory input, such as sound stimulation, promote the transition from sleep to wakefulness (Hayat et al., 2020). Released NA enhances arousal through the action of β and α1 adrenaline receptors located within several subcortical structures, including the medial septal and the medial preoptic area (Berridge, 2008). In this study, the imaging of MS images showed maked increase in NA in the LPO of the hypothalamus immediately after FLs administration (Fig. 6A). The LPO is known to play an important role in regulating the sleep-wake cycle. There are two major cell populations in the LPO: excitatory glutamatergic neurons and inhibitory GABAergic neurons (Taksokhan & Kim, 2023). During sleep, glutamatergic neurons in the LPO are inactive, and GABAergic neurons are active. It is well established that NA projected to the LPO by activation of the LC-NA system inhibits the activity of GABAergic neurons and induces wakefulness (Gallopin et al., 2000; Gallopin et al., 2005; Sangare et al., 2016). The results of this study showed that the FLs-treated group showed a significant increase in spontaneous activity and grooming and rearing, which are indicators of wakefulness, compared with the DW group (Fig. 1). Therefore, it is believed that the LC-NA system was activated after a single oral administration of FL, resulting in wakefulness.

NA neurons from the LC exhibit heightened activity in response to novel stimuli and increased attention, playing a crucial role in arousal-dependent learning and memory (Sara, 2009; Sara & Bouret, 2012). This study observed the impacts on cognition after a single oral dose of FLs using the novel object test (Fig. 2). Mice that underwent memory training one hour after FL administration showed a significant improvement in novel object exploration time and a significant increase in DI. Pharmacological activation of the LC-NA in rodents has been shown to enhance attention and facilitate the shifting of responses from familiar to novel stimuli (Devauges & Sara, 1990). Many episodic-like memories formed in the hippocampus are retained for long periods by a memory stabilization process called early memory fixing (Takeuchi et al., 2016; Yamasaki & Takeuchi, 2017). In the initial stages of memory formation, the release of monoamine from the LC to the hippocampus facilitates the encoding of environmental stimuli and enhances memory retention over a broad time frame of approximately one hour. The findings indicated that FL augmented short-term memory by enhancing LC-dependent early memory consolidation processes, as evidenced by increased novel object exploration time and a significant elevation in DI in mice.

A considerable body of research substantiates that the LC plays a pivotal role in memory formation. For instance, the depletion of monoamines in the LC, the antagonism of NA and DA in multiple brain regions, and the direct inhibition of the LC have all been demonstrated to impair memory across various tasks(Giustino & Maren, 2018; Lisman & Grace, 2005; Uematsu et al., 2017; Wagatsuma et al., 2018). Furthermore, it is established that the activation of the LC and the administration of DA and NA agonists to the LC have a positive effect on memory (Kempadoo et al., 2016; Packard & White, 1989, 1991). Although the LC is believed to regulate neuronal activity by releasing NA, it has been shown that the LC also released dopamine to the hippocampus and other cortical regions (Gálvez-Márquez et al., 2022; Kempadoo et al., 2016). In this experiment, the administration of FLs could not detect projections from the LC to the hippocampus for DA or NA. However, analysis by ISH revealed that the expression of TH, a DA-producing enzyme, and monoamine transporters VMAT2 in LC were induced immediately after the administration of FLs (Fig. 7). There was also a trend towards increased mRNA expression of TH and VMAT2 in the VTA after FL administration. In response to critical events, dopamine neurons in the VTA are known to fire and innervate the CA1 pyramidal layer of the dorsal hippocampus, thereby enhancing long-term memory (Sayegh et al., 2024). Therefore, the series of these results indicate that FLs may stimulate the production of NA and DA in the LC and VTA, which subsequently projects to the hippocampus and enhances short-term memory. However, further research is required to substantiate this hypothesis.

Prior research has indicated that the oral ingestion of FLs may potentially elevate SNS activity. It has been demonstrated that when healthy subjects consumed food rich in FLs, the level of FMD, which signifies an augmentation in peripheral blood flow, exhibited a notable increase two hours later(Hooper et al., 2012; Sun et al., 2019). In our experiments utilizing rodents, a significant enhancement in skeletal muscle blood flow was discerned after the administration of FLs, but an adrenergic receptor inhibitor canceled this change (Saito et al., 2016). This finding suggests that increased blood flow observed after FL ingestion is due to NA being released from sympathetic nerve terminals. CA is released from nerve terminals in response to increased SNS activity and is subsequently secreted into the bloodstream by the adrenal medulla (Osakabe et al., 2023). In mice subjected to stress, the excretion of CA from the blood into the urine indicates SNS activity (Muta et al., 2023). As shown in Fig. 3A, urinary CA concentrations were markedly elevated dose-responsibly. This outcome confirmed that a single dose of FLs elicits activation of SNS. It is well established that increased SNS activity is one of the physiological stress responses and simultaneously activates the HPA axis. Classically, CRH, the master regulator of the HPA axis, has been considered to be secreted from the paraventricular nucleus of the hypothalamus to the pituitary gland in response to stress and to induce adaptation to stress through the release of ACTH and glucocorticoids (Osakabe et al., 2024). Recently, it has been demonstrated that stress stimulation causes the release of CRH from afferents in the hypothalamic paraventricular nucleus, not only into the anterior pituitary gland (Takeuchi et al., 2016) but also into the LC region where CRH receptor 1 is expressed (Aston-Jones et al., 1991; Hauger et al., 2006; Valentino et al., 1983). LC neurons are consequently depolarized, releasing NA from their axon terminals throughout the neuraxis (McCall et al., 2015; Ross & Van Bockstaele, 2020). In this experiment, following administration of FL, CRH mRNA expression was significantly increased in the paraventricular nucleus of the hypothalamus, as shown in Fig. 2B and c. Based on this finding, it is considered that FLs administration acted as a stressor, promoting the release of NA from CRH and LC.

On the other hand, the bioavailability of orally ingested FLs has been demonstrated to be exceedingly low. The probability of achieving effective concentrations in the blood or brain is low(Osakabe et al., 2022; Osakabe & Terao, 2018). Accordingly, the target organ of FL is thought to be the gastrointestinal tract. Previous research has concentrated on alterations in the intestinal microbiota and the secondary metabolites produced in the colon following repeated administration of FLs (Corrêa et al., 2019; Wang et al., 2022). Nevertheless, as the present study demonstrates, a single oral dose of FLs has been observed to alter neurotransmitters in the brain, induce stress responses, and significantly modify mouse behavior immediately. Consequently, it is essential to consider alternative mechanisms.

FLs are a group of substances that have a potent astringent taste and have an essential influence on the sensory properties of chocolate and red wine, affecting their palatability (Ferrer-Gallego et al., 2015; Soares et al., 2020). Astringency is known to polyphenol-specific perception, but the mechanism by which mammals recognize astringent stimuli is currently unknown. Astringency, like other stressors such as capsaicin, may be perceived as a somatosensory (Mouritsen, 2016; Schöbel et al., 2014). According to the previous study, the sensation of astringency remained intact when the gustatory nerve was blocked by local anesthesia but was eliminated when both the trigeminal and gustatory nerves were blocked (Schöbel et al., 2014). In subsequent fMRI studies, astringency stimuli activated several brain areas, including the primary gustatory cortex, which is the insula, superior orbitofrontal cortex, cingulate cortex, and frontal inferior triangular. Besides, this area’s activation intensity was more remarkable than sweet or capsaicin (Kishi et al., 2017). It has been investigated those salient sensory stimuli, including touch, vision, olfaction, and somatosensorial, are input via the paragigantocellular nucleus (PGi), which are located in the anterior medulla oblongata, strongly activates the LC and NTS (Aston-Jones et al., 1991). We observed to increase the NA in the NAc over time following a oral administration of FLs (Fig.6C). It was reported that the NAc receives noradrenergic input from the NTS and LC (Manz et al., 2021). The NTS synthesizes NA, which is released from NTS terminals and enhances memory representations by acting on α-noradrenergic receptors in the NAc (Kerfoot & Williams, 2011). Furthermore, it has been demonstrated that high-fidelity visceral sensory information reaches the NAc via the NTS (McDougall et al., 2024). In light of these findings, it can be reasonably hypothesized that the NA observed in the NAc after FL administration indicates the information transmitted via the NTS as a consequence of stimulation in the gastrointestinal tract.

Astringency of polyphenols including FLs was previously hypothesized to be caused by oral friction caused by the interaction of salivary proline-rich proteins and polyphenols which were detected by mechanoreceptors(Ferrer-Gallego et al., 2015). Conversely, recent studies indicate that salivary proteins may not be essential for perceiving astringency. Recently, it has been proposed that astringency is caused by the activation of transient receptor potential (TRPs) expressed in sensory neurons that sense pungency and reactive oxygen species (ROS). (Kurogi et al., 2015; Takahashi et al., 2021). ROS react with cysteine residues in the ankyrin repeats of TRP channels, causing sensory neurons to fire and transmit stimuli to central nervous system (CNS) (Kozai et al., 2014). Polyphenols with astringency, such as FLs and anthocyanins, are less stable and subject to decomposition by oxidation. This process releases ROS in the neutral pH of the oral cavity and small intestine. It has been documented that this reaction yields low-molecular-weight decomposition products and high-molecular-weight oxides, which are formed by condensation of the decomposition products (Friedman & Jürgens, 2000; Miller et al., 2008; Xue et al., 2024).In our previous study, the increase in skeletal muscle blood flow via hyperactivation of the SNS observed immediately after administration of epicatechin tetramer, a type of flavanol, to rodents was diminished by co-administration of the antioxidant N-acetylcysteine. In addition, this hemodynamic alteration of FL was marked suppressed by co-administration of a TRP-vanilid 1 or TRP-ankirin 1 antagonist (Fushimi et al., 2023). Based on these findings, it is undeniable that ROS produced by certain polyphenols in the gut are perceived as astringency and that this stimulation enhances neurobehavioural and sympathetically mediated circulation and metabolism. Further research is needed to explain how the somatic sensation of astringency is expressed through receptors such as TRP channels.

In the present experiment, alterations in neurotransmitters within the brain were observed following the ingestion of FL. However, several aspects require further elucidation. A notable challenge pertains to the elucidation of the mechanism through which astringent taste is perceived within the oral cavity and digestive tract. Amongst polyphenols, a limited number of compounds that yield astringent stimulation are recognized, and it is hypothesized that the perception of astringent taste is contingent on its chemical structure(Osakabe et al., 2024). As previously mentioned, it is imperative to substantiate the “astringent active oxygen hypothesis” proposing that astringent polyphenols generate ROS and activate TRP channels expressed on gastrointestinal sensory nerves. This can be demonstrated through the utilization of TRP channel knockout mice or electrochemically inactive astringent polyphenol derivatives.

Additionally, it was observed that LC was promptly activated following the intragastric administration of astringent FL. However, the mechanism through which it affects other brain regions remains to be elucidated. In addition to identifying the areas activated by MRI analysis following astringent polyphenol administration, it is essential to visualise the connectivity of each neuron using connectome technology to clarify spatial and temporal changes.

### Conclusion

In conclusion, the results reveal that a single oral dose of astringency FLs promptly activates LC-NA in mice, eliciting stress responses such as hyperactivation of SAM and HPA axis. A series of reactions resulted in the release of NA throughout the brain, significantly impacting attention, alertness, and cognition. Additionally, the activation of memory-related neurons, such as DA neurons in the VTA, was suggested. Moreover, sustained activation of the NAc was observed, leading to the hypothesis that stimulation of the FLs may be perceived as a visceral sensation (Fig.8).

**Fig. 8.**
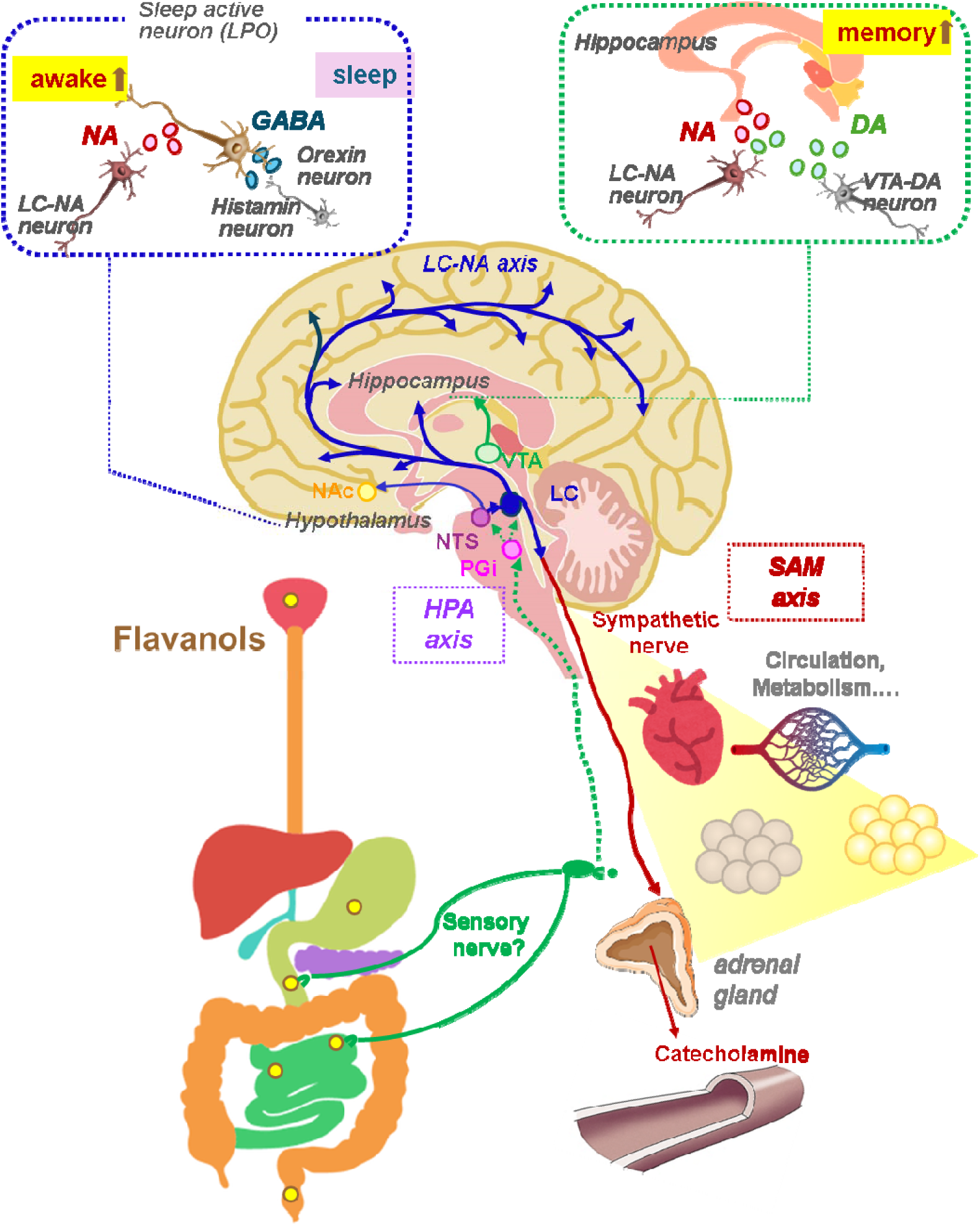
The impact of a single oral administration of flavanols (FLs) on the nervous systems. The oral administration of astringent flavanols (FLs) promptly transmitted a stimulus to the central nervous system, activating the hypothalamic CRH neurons, which subsequently secrete CRH to the locus coeruleus (LC), thereby activating the noradrenaline (NA) neural network. The projection of noradrenaline (NA) from LC to the hypothalamus preoptic area (LPO) has been observed to suppress sleep and promote wakefulness. The projection of NA and dopamine (DA) from LC and DA from the ventral tegmental area (VTA) to the hippocampus was suggested to enhance memory. The projection of NA from LC to the brainstem has been observed to activate sympathetic nerve activity, thereby augmenting circulation and metabolism

However, the precise manner in which FLs’ astringent stimuli are transmitted from the oral cavity and gastrointestinal tract to the brain remains unclear. The provision of a definitive response to this enquiry, and the elucidation of the role played by food sensory properties such as astringency in the preservation of homeostasis, is of significant importance for the maintenance and enhancement of human health.

## Data availability

All data are available in the main text or the supplementary materials.

## Abbreviations

HPA: hypothalamic-pituitary-adrenal
LPO: lateral preoptic area
AD: adrenaline
CA: catecholamine
CNS: central nervous system
CRH: corticotropin-releasing hormone
DI: discrimination index
D.W.: distilled water
DA: dopamine
DBH: dopamine β-hydroxylase
FLs: flavanols
FMD: flow-mediated dilatation
ISH: *in situ* hybridization
LC: locus coeruleus
NA: noradrenalin
NMET: normetanephrine
NAc: nucleus accumbens
PGi: paragigantocellular nucleus
PVN: paraventricular nucleus
ROS: reactive oxygen species
SNS: sympathetic nervous systems
SAM: sympathetic-adreno-medullar
TRP: transient receptor potential
TH: tyrosine hydroxylase
VTA: ventral tegmental area
VMAT: vesicular monoamine transporter

