## Supplemental information for "Astringent flavanol fires the locus-noradrenergic system, regulating neurobehavior and autonomic nerves"

Naomi Osakabe

Department of Bioscience and Engineering, Shibaura Institute of Technology, 307 Fukasaku, Minumaku, Saitama, 337-8570, Japan. Tel +81-48-720-6031, Fax +81-48-720-6011,

### Materials and Methods

#### Materials

(-)-Epicatechin, dimer procyanidin B2, and B5, timer procyanidin C1 and tetramer cinnamtannnin A2 were purchased from Phyto Lab GmbH & Co. KG (Vestenbergsgreuth, ermany, Fig.1a). Caffeine and theobromine were purchased from Sigma-Aldrich (Sigma Aldrich, (St. Louis, MO). The isolation of flavanols (FLs) from cacao liquor was according to the method of Natsume et al. *(17)*. The Prussian Blue method measured total polyphenol concentration in FLs, using (-)- epicatechin as standard. The concentration of each flavanol or xanthin derivatives such as caffeine or theobromine was measured by the method of HPLC *(17)*. The composition of FLs derived from cocoa used in the experiment as follows; (-)-epicatechin (monomer Fig.1a left), 4.56%(W/W); (+)-catechin (monomer Fig.1a left), 6.43%; procyanidin B2(dimer, Fig.1a, right, N=0) , 3.93%; procyanidin C1 (trimer, Fig.1a, right, N=1), 2.36%; cinnamtannin A2 (tetramer, Fig.1a, right, N=2), 1.45 % as shown in Fig.1b.Xanthine derivatives (caffeine, theobromine) were below the detection limit.

#### Animals

10 weeks old male C57BL/6J mice were obtained from CLEA Japan, Inc. (Tokyo, Japan). During two weeks acclimation period, the animals were carefully handled for to reduce anxiety behaviors. Mice housed at room temperature (24-26 °C) under a 12-hour light/dark cycle (light cycle: 7:00-19:00, dark cycle: 19:00-7:00) with free access to water and food. The solid diet (MF) for laboratory animals was obtained from Oriental Yeast Co., Ltd. (Tokyo, Japan). The study was conducted following the Code of Ethics in compliance with the Declaration of Helsinki, and the protocol was approved by the Animal Experimentation Committee of Shibaura Institute of Technology (approval number: AEA 22007). In addition, all animals were humanely raised according to the ARRIVE guidelines of this period.

**Quantification of urinary CA excretion using HPLC**

It has been suggested that brief periods of social isolation in metabolic cages can markedly alter sympathetic nervous system activity in mice *(62).* In our preceding research, we demonstrated that co-housing two mice in metabolic cages markedly reduced the stress response associated with single housing (19). Consequently, in the present study, we employed this approach to examine the sympathetic nervous system activity of FLs. Following a 48-hour acclimation period, urine was collected for 24 hours using a tube containing 20 μl of 2.5 mol/L HCl following oral administration of test chemicals. Oral administration of DW or FLs was performed between 10:00 and 11:00.

**Observation of neurotransmitter location**

Mice were gavaged with either distilled water (DW) or 25 mg/kg flavan 3-ols (FLs) and then decapitated immediately after, 15 or 60 min later. The excised brains were immediately frozen and sectioned (8 μm thick) using a cryostat (Leica, Wetzlar, Germany). The sections of fresh-frozen brains were prepared with a cryostat and thaw-mounted on conductive indium-tin-oxide-coated glass slides (Matsunami Glass Ind. Ltd., Osaka, Japan). A pyrylium-based derivatization method was applied for the tissue localization imaging of NA, its precursors L-dopa and dopamine (DA), and its metabolite normetanephrine (NMET) according to the method by previous publication *(70).* In brief, a solution of TMPy (4.8 mg/200 μL) (Taiyo Nippon Sanso

**Observation of noradrenaline dynamics using ISH**

The mRNA expression of tyrosine hydroxylase (TH), dopamine β-Hydroxylase (DBH), and vesicular monoamine transporter (VMAT) 2 was observed by ISH using consecutive sections from the sections used in the MS imaging analysis. *In situ* hybridization was carried out to evaluate TH, DBH and VMAT mRNA expression with the RNAscope® Multiprex Assay (Advanced Cell Diagnostics) as described above. The target RNA probes were follows; TH (Probe-Mm-Th-C4, 317621-C4, Advanced Cell Diagnostics), DBH (Probe-Mm-Dbh-C3, 400921-C3, Advanced Cell Diagnostics), VMAT2(Probe Mm-Slc18a2, 25331, Advanced Cell Diagnostics).

***Fluorescence microscopy and quantification***

All image analyses were performed by experimenters blinded to the experimental conditions. ISH image was taken using a fluorescence digital microscope (BZ-X800, KEYENCE Corp., Osaka, Japan). Cell counting of c-fos and CRH were performed throughout the extend of PVN. Each expression was quantified using NIH Image J software ver. 1.52 (http://rsb.info.nih.gov/ij/index.html).

**Statistical Analysis**

The sample size was determined from the results of a preliminary experiment using a power test with a significance level of 0.05 and power 0.9. All quantitative assessments were carried out under blinded manner. No data were excluded from the analyses. All data were expressed as mean ± standard deviation. All data were analyzed using GraphPad Prism 10.2.3 (San Diego, CA, USA). The normal distribution of the sample was tested using the Shapiro-Wilk test. Comparisons between the two groups were made using the Student's t test or the Mann-Whitney test. For statistics on urinary CAs, the Kruskal-Wallis test was performed followed by Dunn's multiple comparisons test. Otherwise, two-way ANOVA followed by the Tukey’s multiple comparisons test as post hoc test was used for multiple comparisons. The significance levels were defined at # $p<0.1$, * $p<0.05$, ** $p<0.01$, *** $p<0.001$.

Supplemental results

**Results of spontaneous behavior observation**

The subsequent behavioral trajectories of all mice in the arena over a 120-min period were shown in SFig.1.

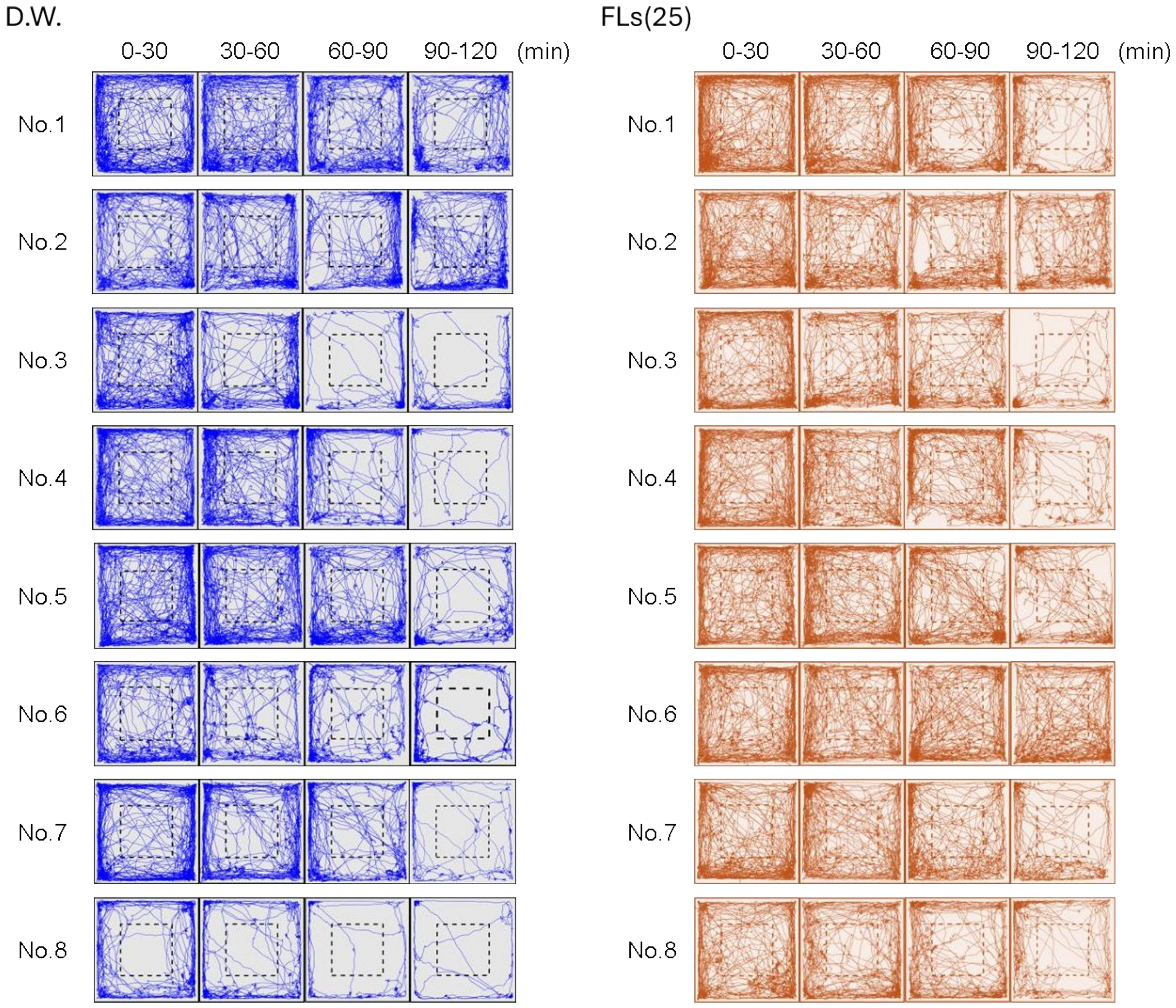


**SFig.1** Behavioral trajectories in the arena for all mice for 120 min following a single oral administration of distilled water or 25mg/kg flavanol (FLs).

**Results of novel object recognition test**

The overall results were shown in Fig. 2. Additionally, typical mouse behavior was also shown in the supplemental video.

**Impact of FLs on HPA axis as observed by mRNA expression of c-fos and CRH in PVN**

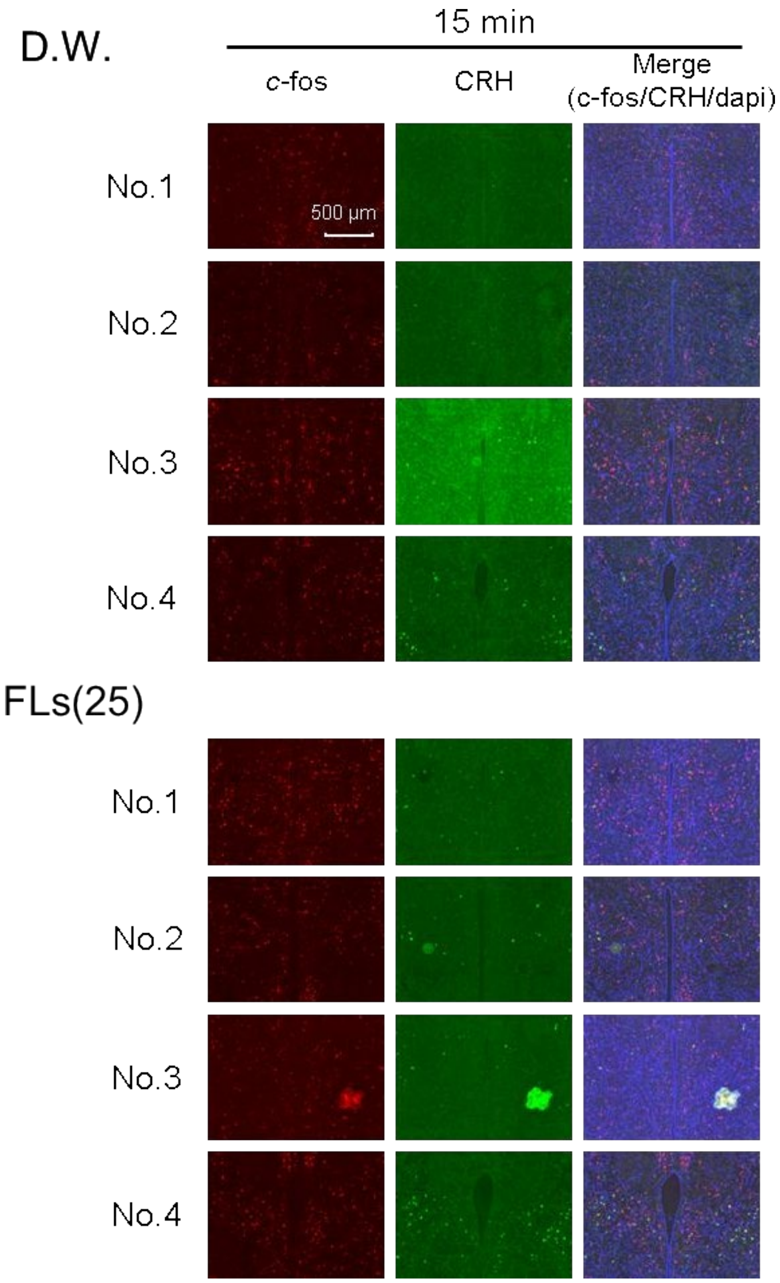

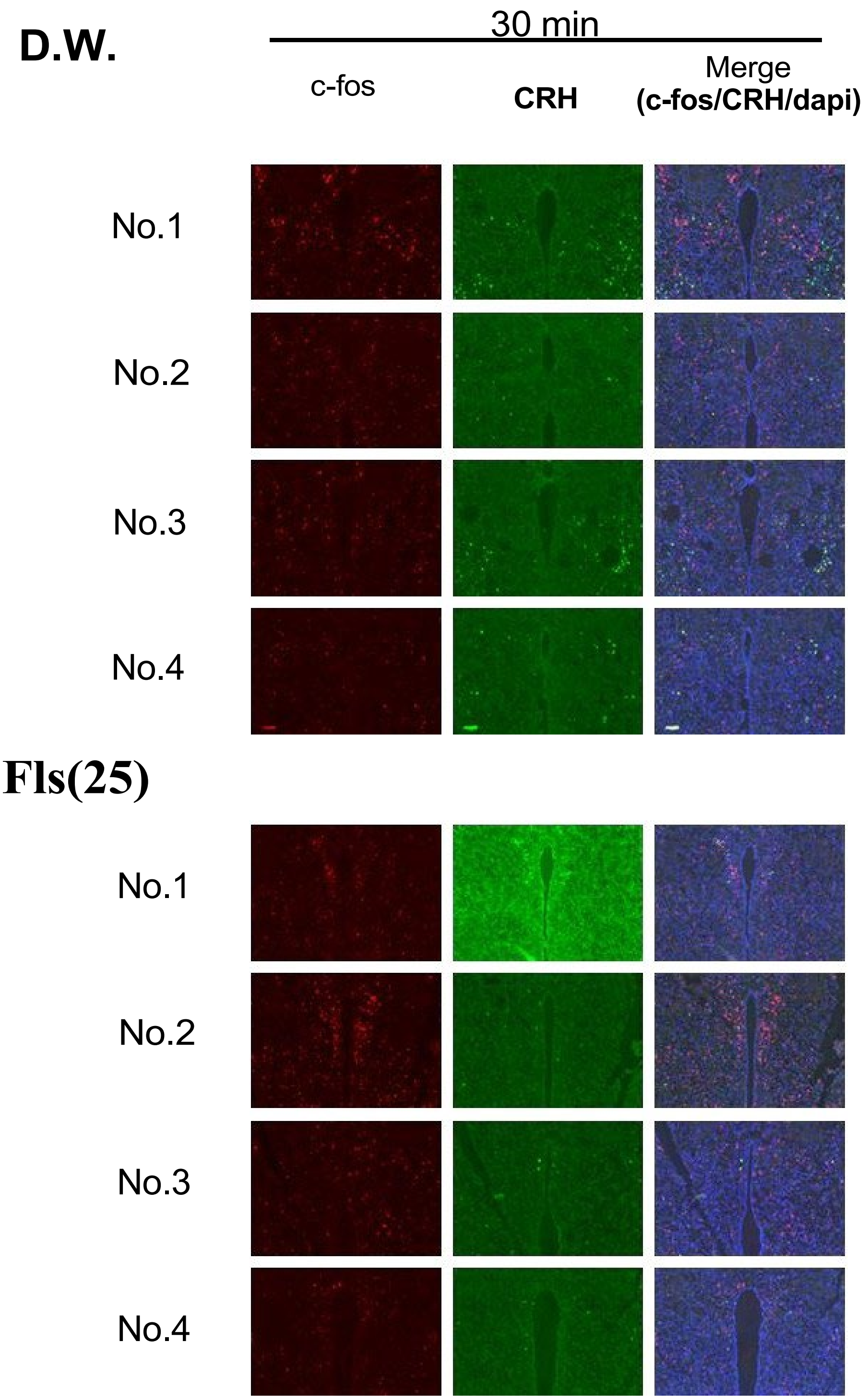
D.W.
30 min
c-fos
CRH
Merge
(c-fos/CRH/dapi)
No.1
No.2
No.3
No.4
Fls(25)
No.1
No.2
No.3
No.4

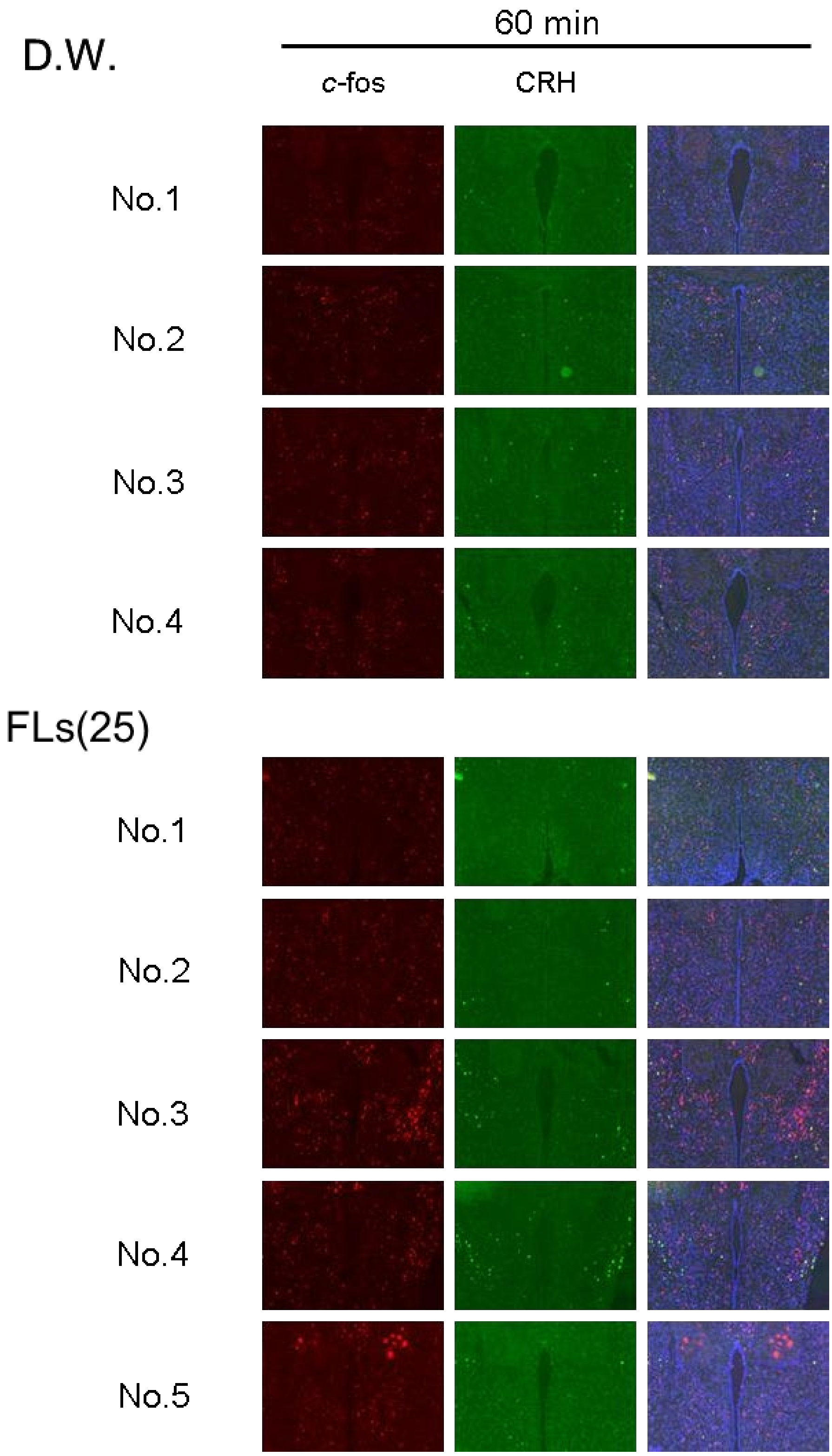


**SFig.2** The results of observing the mRNA expression of cfos and the stress hormone CRH in the hypothalamic PVN of all mice using the ISH method.

**The change of L-dopa, dopamine, noradrenaline and its dynamics in metabolite norethanephrine (NMET) in whole mouse brain observed by the method of MS imaging**

SFig.3 showed the distribution and respective intensities of noradorenaline, its precursors L-dopa and dopamine, and its metabolite norethanephrine detected in sagittal sections of mouse brain immediately after, 15 or 60 min after a single dose of D.W. or 25 mg/kg FL.

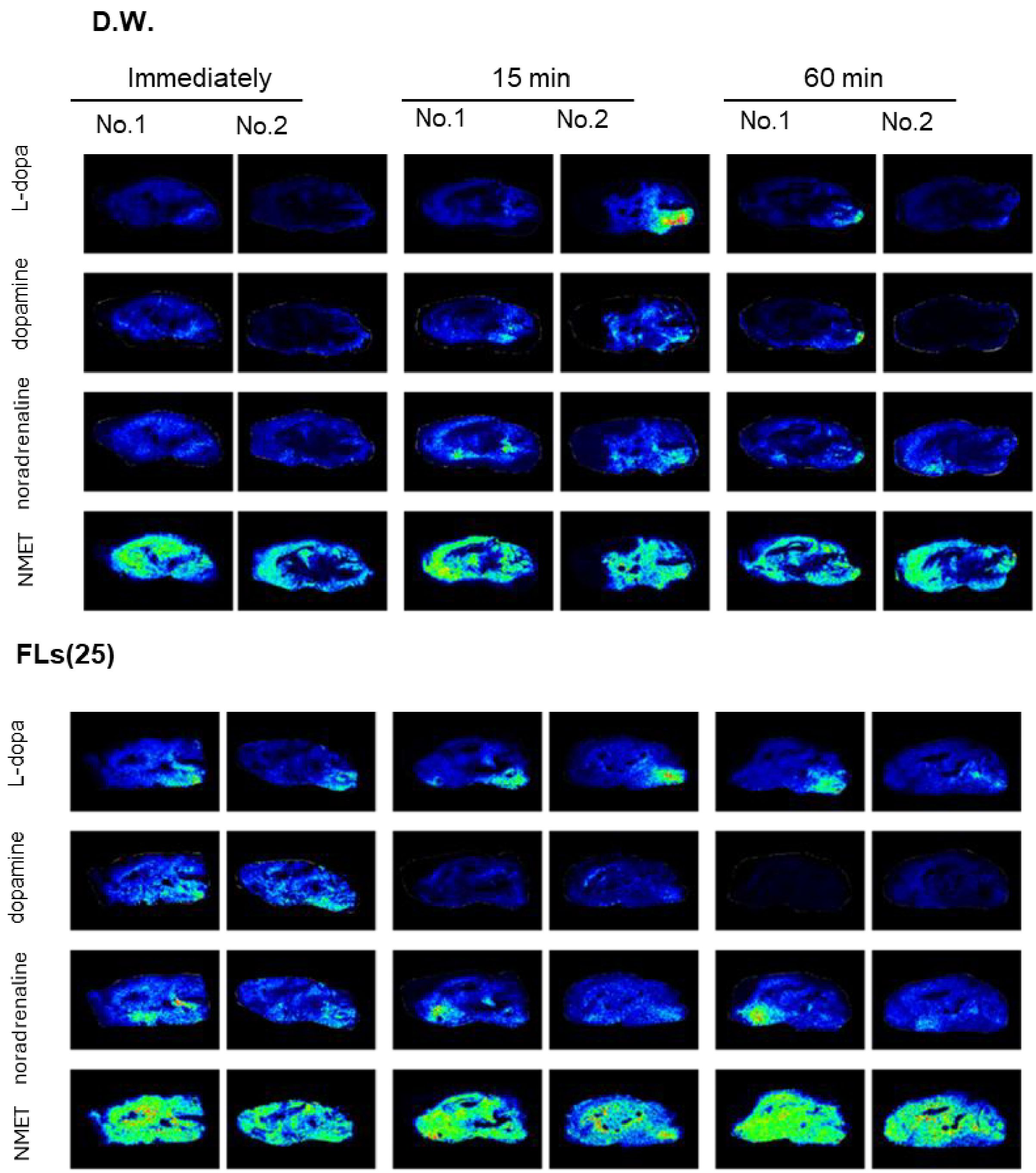

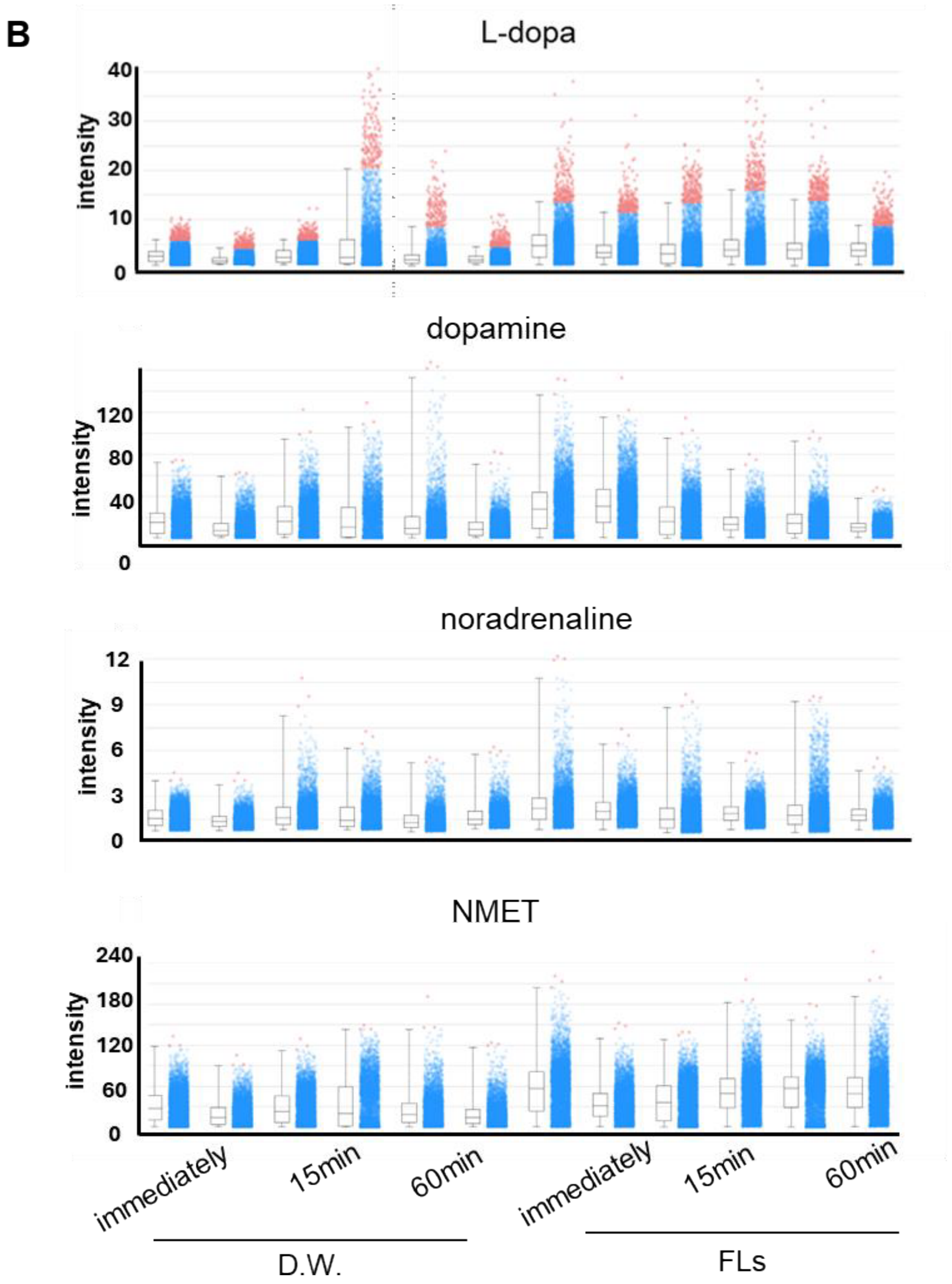


**SFig.3** The intensities of L-dopa, dopamine, noradrenaline and normetanephrine (NMET) immediately, 15 min, and 60 min after a single oral administration of flavanols (FLs) in mice whole brain.A Representative MS images of L-dopa (top), dopamine (DA, second from top), noradrenaline (NA, third from top), and normetanephrine (NMET, bottom). B Relative intensity of L-dopa, DA, NA, and NMET for each animal.

**Intensity of noradrenaline observed in mice LC, LPO, and NAc by the method of MS imaging**

SFig.4 showed the distribution and respective intensities of noradorenaline in LC, LPO, and ACB of mouse brain immediately after, 15 or 60 min after a single dose of D.W. or 25 mg/kg FL.

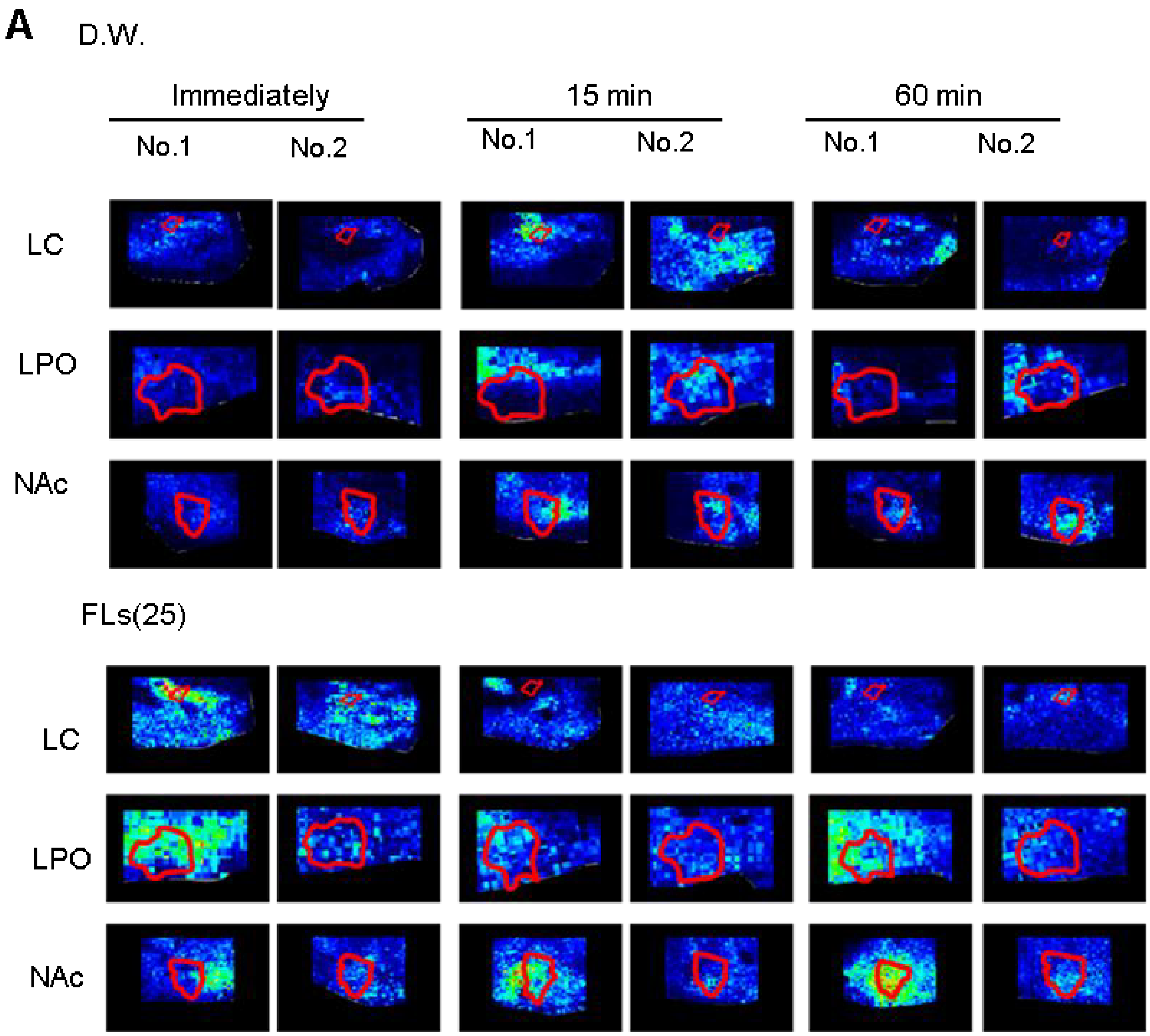

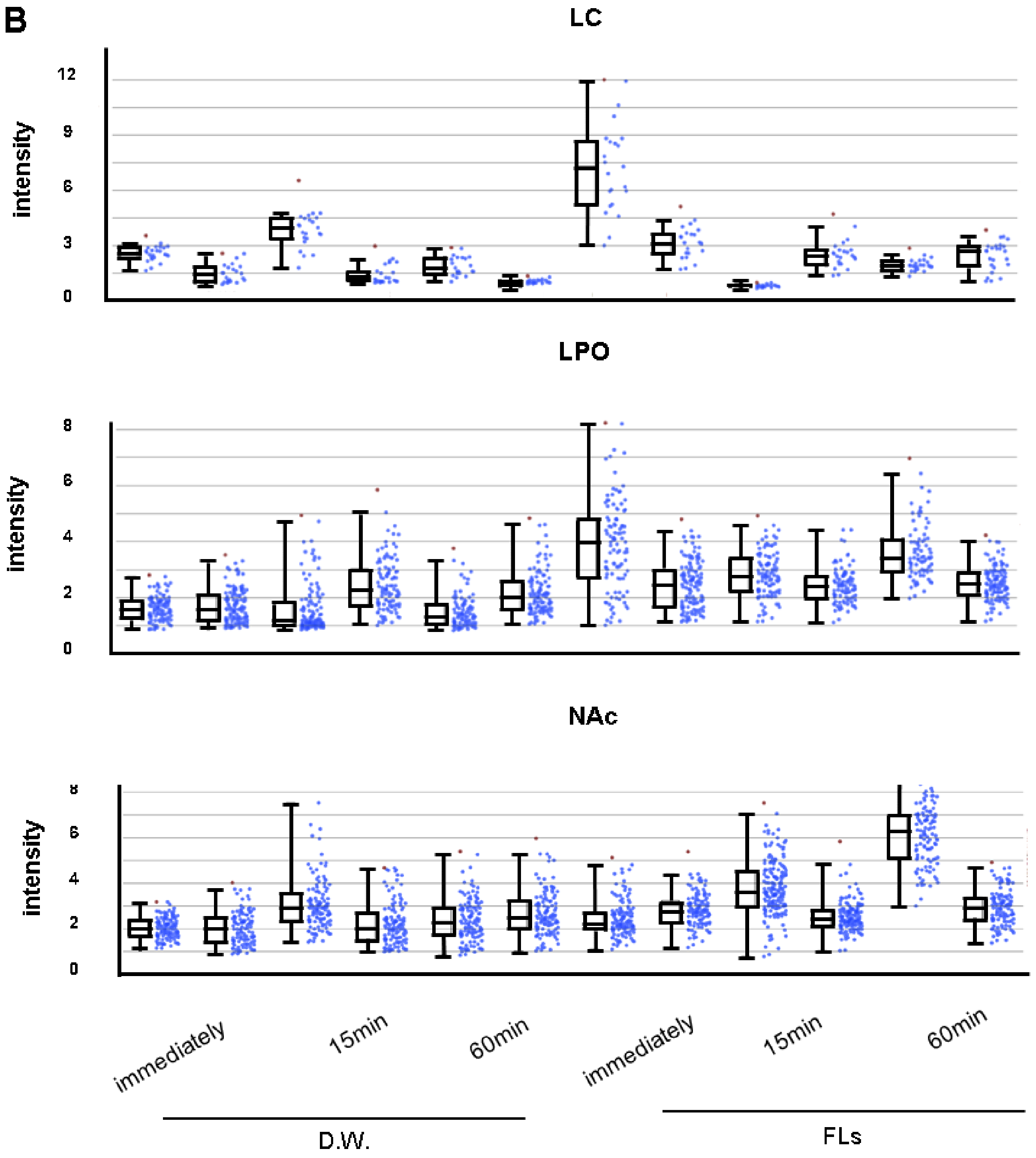


**SFig.4** The intensities of noradrenaline immediately, 15 min, and 60 min after a single oral administration of flavonols (FLs) in locus coeruleus (LC), lateral preoptic area (LPO), and nucleus accumbens (ACB) . A Representative MS images and relative intensity of noradrenaline LC (top), LPO (middle) and ACB( bottom) . B Relative intensity of noradrenalin in LC, LPO and ACB.

**The change of mRNA expression catecholamine synthesis enzymes and its transporter in**

**the brain observed by the method of ISH**

SFig.5 showed the change of mRNA expressions in tyrosine hydroxylase (TH), dopamine β-hydroxylase (DBH), and vesicular monoamine transporter (VMAT)2 in LC and ventral tegmental area (VTA).

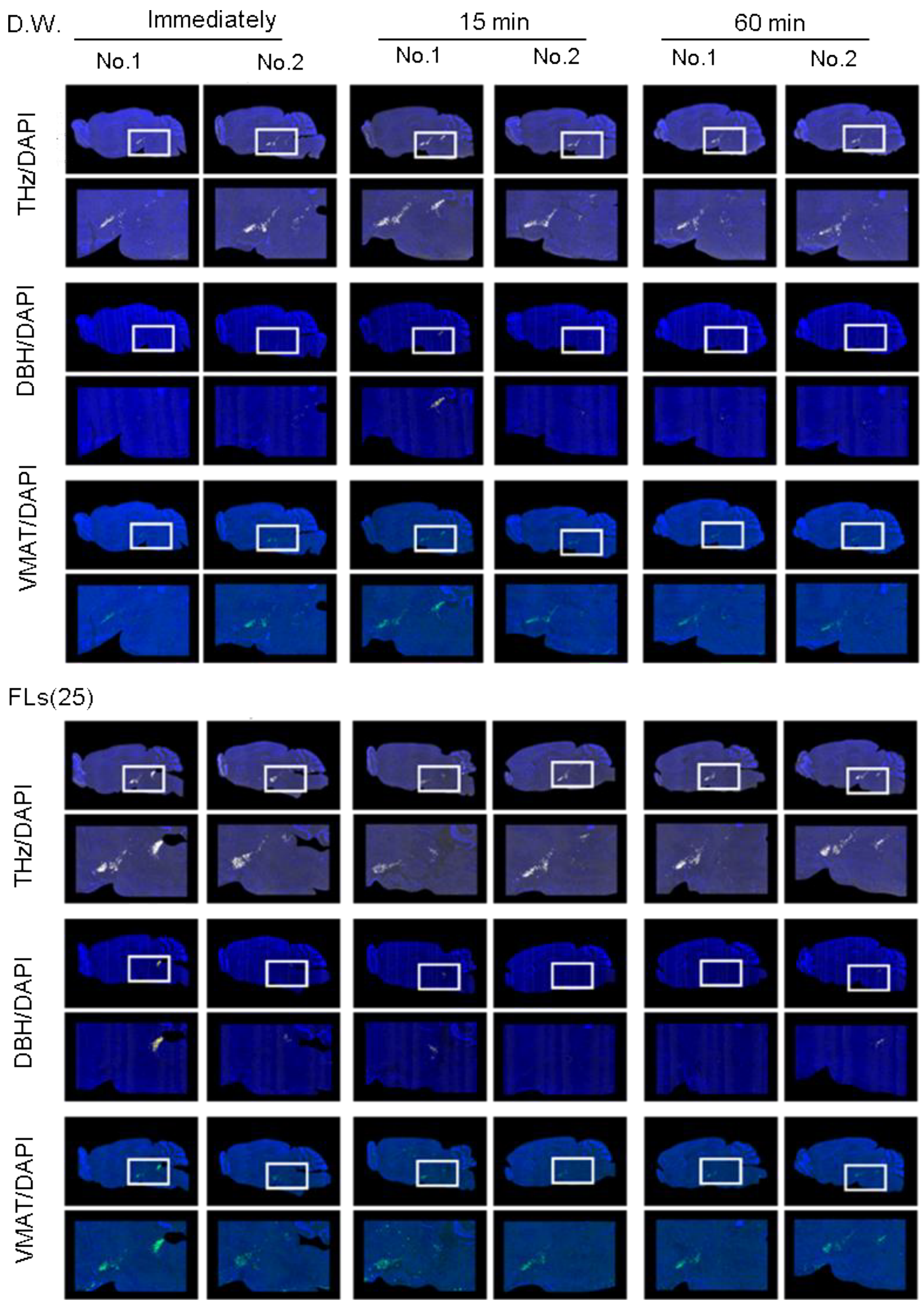

**SFig.5** The change of mRNA expression in TH, DBH and VMAT 2 in LC or VTA of all mice following oral administration of D.W or FLs.

**Supplemental Movie**

Svideo 1
The behavioral trajectory of mice in the NOT test after administration of distilled water(D.W.)
A movie of mouse behavioral trajectory in the NOT test after administration of D.W shown in Fig. 2B(left)

Svideo2
The Behavioral trajectory of mice in the NOT test after administration of Flavanols (FLs)
A movie of mouse behavioral trajectory in the NOT test after administration of FLs shown in Fig. 2B (right)

**Figure 1e**
**Traveled distance every 10min (m)**

| Time (min) | DW | | | | | | | | FLs(25) | | | | | | | |
|---|---|---|---|---|---|---|---|---|---|---|---|---|---|---|---|---|
| | 1 | 2 | 3 | 4 | 5 | 6 | 7 | 8 | 1 | 2 | 3 | 4 | 5 | 6 | 7 | 8 |
| 10 | 30.242 | 25.136 | 32.496 | 21.33 | 33.482 | 23.888 | 31.231 | 19.978 | 50.873 | 40.752 | 33.037 | 32.464 | 29.815 | 38.939 | 31.59 | 32.113 |
| 20 | 28.593 | 18.44 | 25.349 | 30.256 | 34.886 | 21.132 | 31.697 | 18.07 | 37.375 | 28.709 | 27.362 | 28.009 | 36.053 | 26.066 | 26.354 | 19.916 |
| 30 | 28.204 | 22.126 | 23.874 | 29.893 | 25.692 | 23.104 | 26.032 | 13.252 | 28.592 | 27.067 | 29.955 | 32.524 | 22.323 | 30.876 | 32.923 | 26.149 |
| 40 | 27.531 | 20.206 | 23.47 | 27.32 | 30.906 | 15.338 | 20.531 | 15.497 | 32.88 | 22.769 | 26.835 | 24.901 | 34.7 | 24.488 | 28.668 | 25.886 |
| 50 | 25.69 | 21.865 | 19.71 | 27.91 | 27.68 | 17.074 | 26.892 | 13.377 | 25.39 | 26.741 | 22.664 | 25.384 | 31.061 | 24.855 | 22.552 | 22.17 |
| 60 | 27.987 | 21.199 | 13.45 | 21.686 | 24.311 | 16.322 | 20.943 | 12.125 | 25.39 | 26.741 | 22.664 | 25.384 | 31.061 | 24.855 | 22.552 | 22.17 |
| 70 | 24.762 | 19.9 | 12.166 | 26.156 | 28.819 | 13.692 | 21.263 | 9.37 | 30.494 | 21.355 | 21.248 | 25.127 | 30.882 | 31.328 | 25.639 | 24.115 |
| 80 | 21.245 | 18.485 | 8.119 | 8.25 | 24.925 | 14.713 | 18.79 | 4.696 | 29.257 | 25.048 | 22.718 | 22.611 | 24.825 | 28.363 | 23.971 | 23.086 |
| 90 | 21.914 | 17.652 | 6.722 | 15.005 | 15.873 | 16.474 | 15.069 | 8.741 | 18.909 | 20.048 | 18.876 | 25.51 | 21.948 | 29.311 | 18.643 | 14.844 |
| 100 | 17.953 | 22.145 | 8.446 | 8.689 | 13.923 | 8.261 | 15.735 | 9.78 | 24.493 | 21.861 | 13.693 | 15.254 | 17.613 | 28.665 | 14.718 | 19.175 |
| 110 | 18.65 | 20.872 | 13.939 | 6.765 | 17.653 | 17.521 | 7.527 | 4.74 | 13.385 | 19.687 | 8.446 | 10.695 | 18.832 | 28.399 | 18.886 | 14.129 |
| 120 | 14.979 | 20.567 | 3.048 | 1.082 | 6.883 | 5.177 | 5.64 | 1.292 | 10.659 | 19.945 | 6.715 | 10.471 | 14.845 | 25.203 | 11.76 | 13.441 |

**Figure 1f**
**Total travel distance, accumulation (m)**

| Time (min) | DW | | | | | | | | FLs(25) | | | | | | | |
|---|---|---|---|---|---|---|---|---|---|---|---|---|---|---|---|---|
| | 1 | 2 | 3 | 4 | 5 | 6 | 7 | 8 | 1 | 2 | 3 | 4 | 5 | 6 | 7 | 8 |
| 10 | 30.24198 | 25.13628 | 32.49636 | 21.3299 | 33.48205 | 23.88843 | 31.23129 | 19.97831 | 50.87322 | 40.75152 | 33.03747 | 32.46392 | 29.81543 | 38.93866 | 31.59029 | 32.11329 |
| 20 | 58.83535 | 43.57591 | 57.84494 | 51.58607 | 68.36847 | 45.02 | 62.92833 | 38.04819 | 88.24788 | 69.46095 | 60.39898 | 60.47266 | 65.86809 | 65.00508 | 57.94463 | 52.02955 |
| 30 | 87.0389 | 65.70235 | 81.7189 | 81.47881 | 94.06051 | 68.12426 | 88.9603 | 51.29981 | 116.84 | 96.52787 | 90.35436 | 92.99688 | 88.19071 | 95.88076 | 90.86782 | 78.17817 |
| 40 | 114.5695 | 85.90824 | 105.1884 | 108.799 | 124.9669 | 83.4625 | 109.4913 | 66.79658 | 149.7198 | 119.2972 | 117.1898 | 117.8977 | 122.8903 | 120.3692 | 119.5363 | 104.064 |
| 50 | 140.2593 | 107.7734 | 124.8986 | 136.7089 | 152.6471 | 100.5361 | 136.3836 | 80.17357 | 175.1098 | 146.0387 | 139.8534 | 143.2817 | 153.951 | 145.2241 | 142.0881 | 126.2343 |
| 60 | 168.2465 | 128.9724 | 138.349 | 158.3953 | 176.958 | 116.8577 | 157.3263 | 92.29818 | 200.4998 | 172.7801 | 162.5169 | 168.6657 | 185.0117 | 170.0789 | 164.6399 | 148.4045 |
| 70 | 193.0081 | 148.8727 | 150.5152 | 184.5518 | 205.7774 | 130.5493 | 178.5889 | 101.6677 | 230.9939 | 194.1348 | 183.7648 | 193.793 | 215.8936 | 201.4064 | 190.2786 | 172.519 |
| 80 | 214.2535 | 167.3579 | 158.6343 | 192.8019 | 230.7019 | 145.2621 | 197.3787 | 106.3636 | 260.2511 | 219.1829 | 206.4824 | 216.4045 | 240.7186 | 229.7693 | 214.25 | 195.6049 |
| 90 | 236.1673 | 185.0094 | 165.356 | 207.8071 | 246.5752 | 161.7359 | 212.4477 | 115.1046 | 279.1601 | 239.2305 | 225.3588 | 241.9149 | 262.6667 | 259.0801 | 232.8927 | 210.4492 |
| 100 | 254.1199 | 207.1541 | 173.8022 | 216.496 | 260.4986 | 169.9966 | 228.1828 | 124.8848 | 303.6528 | 261.0918 | 239.0521 | 257.1693 | 280.2798 | 287.745 | 247.6111 | 229.6242 |
| 110 | 272.7702 | 228.0264 | 187.7411 | 223.261 | 278.1517 | 187.5174 | 235.7097 | 129.6252 | 317.0382 | 280.7791 | 247.4979 | 267.8647 | 299.1119 | 316.144 | 266.4975 | 243.7528 |
| 120 | 287.7489 | 248.5937 | 190.7895 | 224.343 | 285.035 | 192.6939 | 241.3498 | 130.9168 | 327.6974 | 300.724 | 254.2131 | 278.3356 | 313.9565 | 341.3471 | 278.2576 | 257.1939 |

| | **Figure 1g** **Total travel distance (m)** | | **Figure 1h** **Total spent time in center area (s** | | **Figure 1i** **Grooming(number)** | | **Figure 1j** **Rearing(number)** | |
|---|---|---|---|---|---|---|---|---|
| | DW | FLs(25) | DW | FLs(25) | DW | FLs(25) | DW | FLs(25) |
| 1 | 287.7489 | 327.6974 | 311.16 | 379.12 | 52 | 69 | 383 | 680 |
| 2 | 248.5937 | 300.724 | 407.76 | 842.2 | 42 | 46 | 432 | 584 |
| 3 | 190.7895 | 254.2131 | 143.56 | 358.44 | 33 | 63 | 290 | 479 |
| 4 | 224.343 | 278.3356 | 296 | 453.68 | 22 | 59 | 373 | 470 |
| 5 | 285.035 | 313.9565 | 323.4 | 527.84 | 27 | 62 | 468 | 734 |
| 6 | 192.6939 | 341.3471 | 359.64 | 817.72 | 41 | 46 | 363 | 756 |
| 7 | 241.3498 | 278.2576 | 256.92 | 497.64 | 41 | 56 | 418 | 606 |
| 8 | 130.9168 | 257.1939 | 38.44 | 343.64 | 41 | 40 | 200 | 524 |

**Figure 2c**
**Exploration time (sec)**

**Figure 2d**
**DI**

| ID | DW | | FLs(25) | | DI DW | DI FLs(25) |
|---|---|---|---|---|---|---|
| - | Familiar | Novel | Familiar | Novel | | |
| 1 | 14.875 | 14.29 | 9.995 | 18.21 | 0.489971 | 0.64563 |
| 2 | 7.275 | 11.825 | 10.36 | 22.275 | 0.61911 | 0.682549 |
| 3 | 11.14 | 7.8 | 5.335 | 19.835 | 0.411827 | 0.788041 |
| 4 | 9.925 | 11.235 | 8.715 | 25.375 | 0.530955 | 0.744353 |
| 5 | 12.58 | 12.335 | 10.815 | 19.78 | 0.495083 | 0.646511 |
| 6 | 11.105 | 8.06 | 9.05 | 20.715 | 0.420558 | 0.695952 |
| 7 | 11.51 | 11.35 | 10.525 | 11.135 | 0.4965 | 0.514081 |
| 8 | 9.11 | 11.935 | 8.885 | 20.125 | 0.567118 | 0.693726 |

**Figure 3b**
**Urine adrenaline (ng/μg Cre)**

| ID _ | DW | FLs(25) | FLs(50) |
|---|---|---|---|
| 1 | 0.069 | 0.173 | 0.28 |
| 2 | 0.108 | 0.165 | 0.163 |
| 3 | 0.145 | 0.122 | 0.274 |
| 4 | 0.162 | 0.145 | 0.338 |
| 5 | 0.06 | 0.213 | 0.322 |
| 6 | 0.142 | 0.051 | 0.283 |
| 7 | 0.116 | 0.277 | 0.277 |
| 8 | 0.15 | 0.194 | 0.377 |
| 9 | 0.175 | | |
| 10 | 0.184 | | |
| 11 | 0.185 | | |
| 12 | 0.174 | | |
| 13 | 0.241 | | |
| 14 | 0.228 | | |
| 15 | 0.245 | | |

**Figure 3c**
**Urine noradrenaline (ng/μg Cre)**

| ID | DW | FLs(25) | FLs(50) |
|---|---|---|---|
| 1 | 0.069 | 0.173 | 0.28 |
| 2 | 0.108 | 0.165 | 0.163 |
| 3 | 0.145 | 0.122 | 0.274 |
| 4 | 0.162 | 0.145 | 0.338 |
| 5 | 0.06 | 0.213 | 0.322 |
| 6 | 0.142 | 0.051 | 0.283 |
| 7 | 0.116 | 0.277 | 0.277 |
| 8 | 0.15 | 0.194 | 0.377 |
| 9 | 0.175 | | |
| 10 | 0.184 | | |
| 11 | 0.185 | | |
| 12 | 0.174 | | |
| 13 | 0.241 | | |
| 14 | 0.228 | | |
| 15 | 0.245 | | |

**Figure 3d**
**Urine noradrenaline + adrenaline (ng/μg Cre)**

| ID | DW | FLs(25) | FLs(50) |
|---|---|---|---|
| 1 | 0.37 | 0.383 | 0.843 |
| 2 | 0.356 | 0.529 | 0.613 |
| 3 | 0.416 | 0.521 | 0.781 |
| 4 | 0.555 | 0.473 | 0.731 |
| 5 | 0.321 | 0.559 | 0.813 |
| 6 | 0.275 | 0.398 | 0.847 |
| 7 | 0.406 | 0.701 | 0.791 |
| 8 | 0.602 | 0.34 | 0.874 |
| 9 | 0.585 | | |
| 10 | 0.52 | | |
| 11 | 0.587 | | |
| 12 | 0.506 | | |
| 13 | 0.723 | | |
| 14 | 0.674 | | |
| 15 | 0.569 | | |

**Figure 4c**
**c-fos mRNA labeled cells**

| Time (min) | DW | | | | FLs(25) | | | | |
|---|---|---|---|---|---|---|---|---|---|
| ID | 1 | 2 | 3 | 4 | 1 | 2 | 3 | 4 | 5 |
| 15 | 16 | 7 | 26 | 13 | 25 | 21 | 17 | 18 | |
| 30 | 23 | 12 | 14 | 10 | 35 | 50 | 15 | 9 | |
| 60 | 5 | 23 | 15 | 7 | 9 | 12 | 9 | 6 | 13 |

**Figure 4d**
**CRH mRNA labeled cells**

| Time (min) | DW | | | | FLs(25) | | | | |
|---|---|---|---|---|---|---|---|---|---|
| ID | 1 | 2 | 3 | 4 | 1 | 2 | 3 | 4 | 5 |
| 15 | 8 | 9 | 11 | 5 | 13 | 3 | 10 | 1 | |
| 30 | 4 | 10 | 0 | 2 | 37 | 24 | 7 | 30 | |
| 60 | 6 | 16 | 7 | 3 | 13 | 1 | 0 | 3 | 9 |
